# Fast rule switching and slow rule updating in a perceptual categorization task

**DOI:** 10.1101/2022.01.29.478330

**Authors:** F. Bouchacourt, S. Tafazoli, M.G. Mattar, T.J. Buschman, N.D. Daw

## Abstract

To adapt to a changing world, we must be able to switch between rules already learned and, at other times, learn rules anew. Often we must do both at the same time, switching between known rules while also constantly re-estimating them. Here, we show these two processes, rule switching and rule learning, rely on distinct but intertwined computations, namely fast inference and slower incremental learning. To this end, we studied how monkeys switched between three rules. Each rule was compositional, requiring the animal to discriminate one of two features of a stimulus and then respond with an associated eye movement along one of two different response axes. By modeling behavior we found the animals learned the axis of response using fast inference (*rule switching*) while continuously re-estimating the stimulus-response associations within an axis (*rule learning*). Our results shed light on the computational interactions between rule switching and rule learning, and make testable neural predictions for these interactions.

## Introduction

A crucial component of intelligence is learning from the environment, allowing one to modify their behavior in light of experience. A long tradition of research in areas like Pavlovian and instrumental conditioning has focused on elucidating general learning mechanisms – especially error-driven incremental learning rules associated with dopamine and the basal ganglia^1–10^. More recently, it has become increasingly clear that the brain’s dynamics for learning can themselves be adapted^11,12^. The general-purpose incremental learning mechanisms can allow animals to gradually learn a new stimulus-response discrimination rule by trial and error. But this type of gradual adjustment seems unlikely to account for other more task-specialized learning effects: for instance, if two different stimulus-response rules are repeatedly reinforced in alternation, animals can come to switch between them more rapidly^13–15^. Such rule switching has many formal similarities with *de novo* rule learning: here also, response behavior is modified in light of feedback, often progressively over several trials. However these more task-specialized dynamics are often modeled by a distinct computational mechanism – e.g., Bayesian inference, where animals accumulate evidence about which rule currently applies^16–19^. Inference is associated with activity in the prefrontal cortex^20–33^, suggesting the neural mechanisms are distinct from incremental learning.

This type of inference process presupposes that the animal has previously learned about the structure of the task: which rules can apply, how often they switch, etc., and so it has typically been studied in well-trained animals^12,16,34^. However, there has been increasing theoretical interest – but relatively little direct empirical evidence – in the mechanisms by which the brain learns the broader structure of the task in order to build task-specialized inference mechanisms for rapid rule switching. For Bayesian inference models, this problem corresponds to learning the generative model of the task, e.g. inferring a mixture model over latent contexts or states (rules or task conditions, which are ‘latent’ since they are not overtly signaled) and their properties (e.g., stimulus-response-reward contingencies)^16,17,35–40^. A more mechanistic, but not mutually exclusive, account of rule switching suggests it may be implemented by the dynamics of a recurrent neural network (RNN), in which case higher-level learning (here called ‘metalearning’) corresponds to tuning the weights of this network^11,39,41–44^.

Perhaps most intriguingly, these accounts often posit that a full theory of rule learning and rule switching ultimately involves an interaction between both major classes of learning mechanisms, inferential and incremental. Thus, in Bayesian inference models, it is often hypothesized that an inferential process (e.g. prefrontal) decides which latent state is in effect, while the properties of each state are learned, conditional on this, by downstream (e.g. striatal) error-driven incremental learning^17,39^. Of course, this conventional distinction between “inferential” rule switching over “incremental” learned rules is imperfect: inference about the world’s current state can be slow (e.g., to the extent evidence accumulates gradually) and learning about the properties of these states can itself be understood as inference, at a different level of a hierarchical model. Somewhat similarly, metalearning of RNN dynamics for rule switching has been proposed to be itself driven by incremental error-driven updates^7,11,42^. However, apart from a few interesting examples in human rule learning^17,35,45–48^, this type of interaction has mostly been posited theoretically, while the two learning mechanisms have mostly been studied in regimes where they operate more or less in isolation^12,16,34,49–53^.

To study rule switching and rule learning, we trained non-human primates to perform a rule-based category-response task. Depending on the rule in effect, the animals needed to attend to and categorize either the color or the shape of a stimulus, and then respond with a saccade along one of two different response axes. We observe a combination of both fast and slow learning during the task: monkeys rapidly switched into the correct response axis, consistent with inferential learning of the response state, while, within a state, the animals slowly learned category-response mappings, consistent with incremental (re)learning, and even though they were well-trained on the task beforehand. To quantify the learning mechanisms underlying the animals’ behavior, we tested whether inference or incremental classes of models, separately, could explain the behavior. Both classes of models reproduced learning-like effects – i.e., dynamic, experience-driven changes in behavior. However, neither model could, by itself, explain the combination of both fast and slow learning. Importantly, the fact that a fully informed inferential learner could not explain the behavior also indicated that the observed fast and slow learning was not simply driven by informed adjustment to the task structure itself. Instead, we found that key features of behavior were well explained by a hybrid rule-switching and rule-learning model, which inferred which response axis was active while continually performing slower, incremental relearning of the consequent stimulus-response mappings within an axis. These results support the hypothesis that there are multiple, interacting, mechanisms that guide behavior in a contextually-appropriate manner.

## Results

### Task design and performance

Two rhesus macaques were trained to perform a rule-based category-response task. On each trial, the monkeys were presented with a stimulus that was composed of a color and shape (Fig. 1a). The shape and color of each stimulus were drawn from a continuous space and, depending on the current rule in effect, the animals categorized the stimulus according to either its color (red vs. green) or its shape (‘bunny’ vs.‘tee’, Fig. 1b). Then, as a function of the category of the stimulus and the current rule, the animals made one of four different responses (an upper left, upper right, lower left, or lower right saccade).

**Figure 1:**
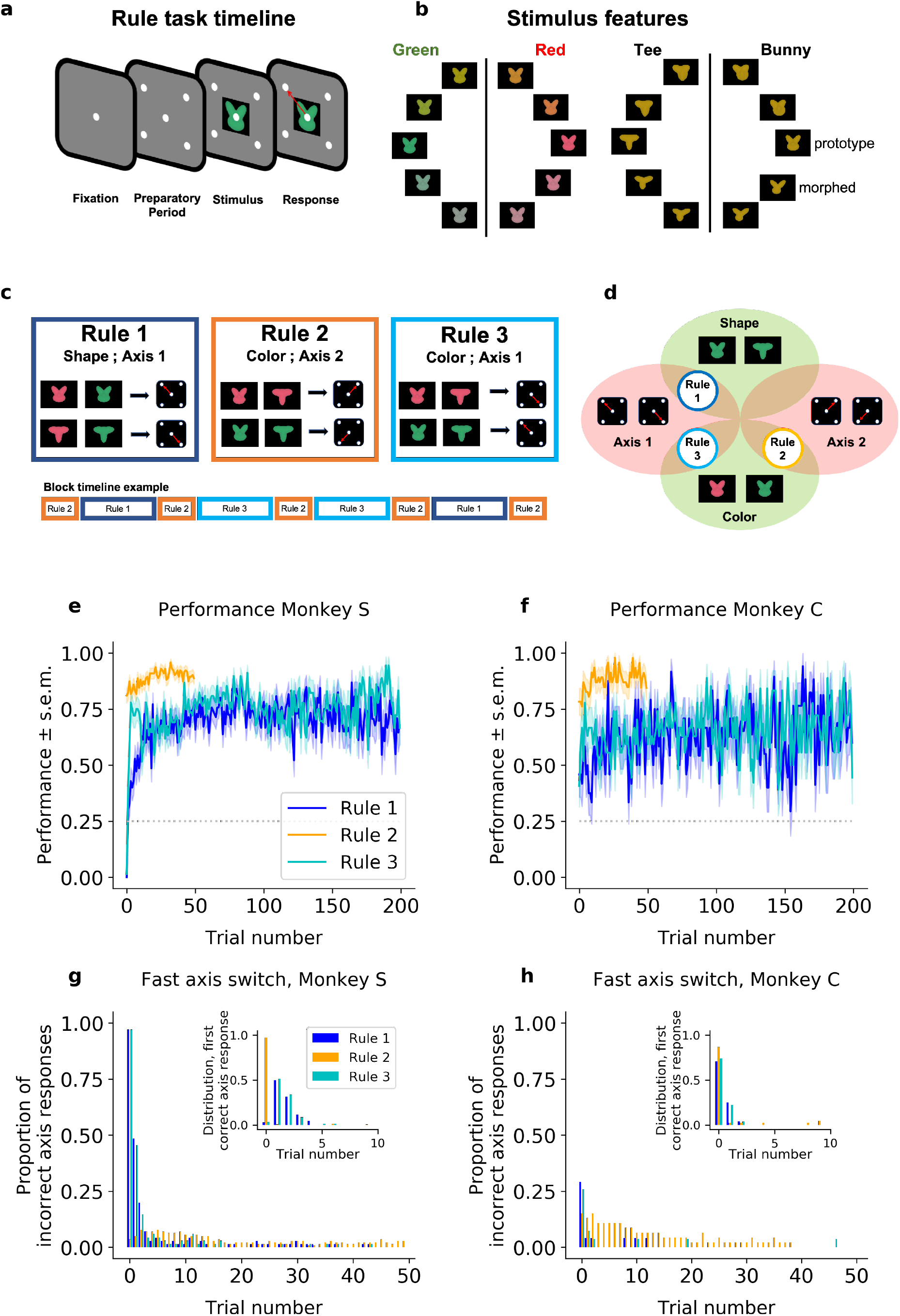
Task design and performance (including all trials). (a) Schematic of a trial. (b) Stimuli were drawn from a two-dimensional feature space, morphing both color (left) and shape (right). Stimulus categories are indicated by vertical lines and labels. (c) The stimulus-response mapping for the three rules, and an example of a block timeline. (d) Venn diagram showing the overlap between rules. (e,f) Average performance (sample mean and standard error of the mean) for each rule, for (e) Monkey S and (f) Monkey C. (g,h) Proportion of responses on the incorrect axis for the first 50 trials of each block for (g) Monkey S and (h) Monkey C. Insets: Trial number of the first response on the correct axis after a block switch, respectively for Monkey S and C.

Animals were trained on three different category-response rules (Fig. 1c). Rule 1 required the animal to categorize the shape of the stimulus, making a saccade to the upper-left location when the shape was categorized as a ‘bunny’ and a saccade to the lower-right location when the shape was categorized as a ‘tee’. These two locations – upper-left and lower-right – formed an ‘axis’ of response (*Axis 1*). Rule 2 was similar but required the animal to categorize the color of the stimulus and then respond on the opposite axis (*Axis 2;* red=upper-right, green=lower-left). Finally, Rule 3 required categorizing the color of the stimulus and responding on *Axis 1* (red=lower-right, green=upper-left). Note that these rules are compositional in nature, with overlapping dimensions (Fig. 1d). Rule 1 required categorizing the shape of the stimulus, while Rules 2 and 3 required categorizing the color of the stimulus. Similarly, Rules 1 and 3 required responding on the same axis (*Axis 1*), while Rule 2 required a different set of responses (*Axis 2*). In addition, the overlap in the response axis for Rules 1 and 3 meant certain stimuli had congruent responses for both rules (e.g., red-tee and green-bunny stimuli) while other stimuli had incongruent responses between rules (e.g., red-bunny and green-tee). For all rules, when the animal made a correct response, it received a reward (an incorrect response led to a short ‘time-out’).

Animals performed the same rule during a block of trials. Critically, the animals were not explicitly cued as to which rule was in effect for that block. Instead, they had to use information about the stimulus, their response, and reward feedback, to infer which rule was in effect. After the animals discovered the rule and were performing it at a high level (defined as >70%, see Methods) the rule would switch. Although unpredictable, the moment of switching rules was cued to the animal (with a flashing screen). Importantly, this switch-cue did not indicate which rule was now in effect (just that a switch had occurred). To facilitate learning and performance, the sequence of rules across blocks was semi-structured such that the axis of response always changed following a block switch (i.e. after a Rule 2 block the animal performed either Rule 1 or Rule 3, chosen pseudo randomly, and vice versa, see block timeline example in Fig. 1c). Note that the cue implied an axis switch but did not instruct which axis to use, thus the animals must learn and continually track on which axis to respond.

Overall, both monkeys performed the task well above chance (Fig. 1e,f). When the rule switched to Rule 2, the animals quickly switched their behavior: Monkey S responded correctly on the first trial in 81%, CI=[0.74,0.87] of Rule 2 blocks, and reached 91%, CI=[0.85,0.95] after only 20 trials (Monkey C being respectively at 78%, CI=[0.65,0.88]; and 85%, CI=[0.72,0.92]). In Rule 1 and Rule 3, their performance also exceeded chance level quickly. In Rule 1, although the performance of Monkey S was below chance on the first trial (0%, CI=[0,0.052]; 46%, CI=[0.28,0.65] for Monkey C), reflecting perseveration on the previous rule, performance quickly climbed above chance (77% after 50 trials, CI=[0.66,0.85]; 63%, CI=[0.43,0.79] for Monkey C). A similar pattern was seen for Rule 3 (initial performance of 1.5%, CI=[0.0026,0.079] and 78%, CI=[0.67,0.86] after 50 trials for Monkey S; 41%, CI=[0.25,0.59] and 67%, CI=[0.48,0.81] for Monkey C, respectively).

While the monkeys performed all three rules well, there were two interesting behavioral phenomena. First, the monkeys were slower to switch to Rule 1 and Rule 3 than to switch to Rule 2. On the first 20 trials, the difference in average percent performance of Monkey S was Δ=35 between Rule 2 and Rule 1, and Δ=22 between Rule 2 and Rule 3 (significant Fisher test comparing Rule 2 to Rule 1 and Rule 3, with p<10^(−4)^ in both conditions; respectively Δ=35, Δ=24 and p<10^(− 4)^ for Monkey C).

Second, both monkeys learned the axis of response nearly instantaneously. After a switch cue, Monkey S almost always responded on Axis 2 (the response axis consistent with Rule 2; 97%, CI=[0.90,0.99] in Rule 1; 97%, CI=[0.92,0.98] in Rule 2; 97%, CI=[0.90,0.99] in Rule 3; see Fig. 1g). Then, if this was incorrect, it switched to the correct axis within 5 trials on 97%, CI=[0.90,0.99] of blocks of Rule 1, and 94% CI=[0.86,0.98] of blocks of Rule 3. Monkey C instead tended to alternate the response axis on the first trial following a switch cue (it made a response on the correct axis on the first trial with a probability of 71%, CI=[0.51,0.85] in Rule 1; 85%, CI=[0.72,0.92] in Rule 2; and 84%, CI=[0.55,0.87] in Rule 3), implying an understanding of the pattern of axis changes with block switches (Fig. 1h). Both monkeys maintained the correct axis with very few off-axis responses throughout the block (at trial 20, Monkey S: 1.4%, CI=[0.0025,0.077] in Rule 1; 2.1%, CI=[0.0072,0.060] in Rule 2; 0%, CI=[0,0.053] in Rule 3; Monkey C: 0%, CI=[0,0.14] in Rule 1; 4.3%, CI=[0.012,0.15] in Rule 2; 3.7%, CI=[0.0066,0.18] in Rule 3). These results suggest the animals were able to quickly identify the axis of response but took longer (particularly for Rules 1 and 3) to learn the correct mapping between stimulus features and responses within an axis.

### Learning rules *de novo* cannot capture the behavior

To perform the task, the animals had to learn which rule was in effect during each block of trials. This required determining both the response axis and the relevant feature. As noted above, the monkeys’ behavior suggests learning was a mixture of fast switching, reminiscent of inference models, and slow refinement, as in error-driven incremental learning. Given this, we began by testing whether the inference or incremental classes of models could capture the animals’ behavior. As in previous work^54–58^, all our models shared common noisy perceptual input and action selection stages (Fig. S1, and Methods). As we detail next, the intervening mechanism for mapping stimulus to action value differed between models.

First, we fit an error-driven learning model, which gradually relearns the stimulus-response mappings *de novo* at the start of each block, to the monkeys’ behavior. This class of models works by learning the reward expected for different stimulus-response combinations, using incremental running averages to smooth out trial-to-trial stochasticity in reward realization – here, due to perceptual noise in the stimulus classification. In particular, we fit a variant of Q learning (model QL, see Fig. S1 and Methods) that was elaborated to improve its performance in this task: for each action, model QL parameterized the mapping from stimulus to reward linearly using two basis functions over the feature space (one binary indicator each for color and shape), and used error-driven learning to estimate the appropriate weights on these for each block. This scheme effectively builds-in the two relevant feature-classification rules (shape and color), making the generous assumption that the animals had already learned the categories. In addition, the model resets the weights to fixed initial values at each block switch, allowing the model to start afresh and avoiding the need to unlearn. Yet, even with these built-in advantages, the model was unable to match the animals’ ability to rapidly switch axes, but instead relearned the feature-response associations after each block switch (simulations under best-fitting parameters shown in Fig. 2 for Monkey S, Fig. S2 for Monkey C). Several tests verified the model learned more slowly than the animals (Fig. 2a,b). For instance, the model fitted on Monkey S’s behavior responded on Axis 2 on the first trial of the block only 50% of the time in all three rules (Fig. 2a, Fisher test of model simulations against data: p<10^(−4)^ for the three rules). The model thus failed to capture the initial bias of Monkey S for Rule 2 discussed above. Importantly, the model switched to the correct axis within 5 trials on only 58% of blocks of Rule 1, and 57% of blocks of Rule 3 (Fig. 2b, Fisher test against monkey behavior: p<10^(−4)^). Finally, the model performed 24% of off-axis responses after 20 trials in Rule 1, 21% in Rule 2, and 21% in Rule 3, all much higher than what was observed in the monkey’s behavior (Fisher test p<10^(−4)^).

**Figure 2:**
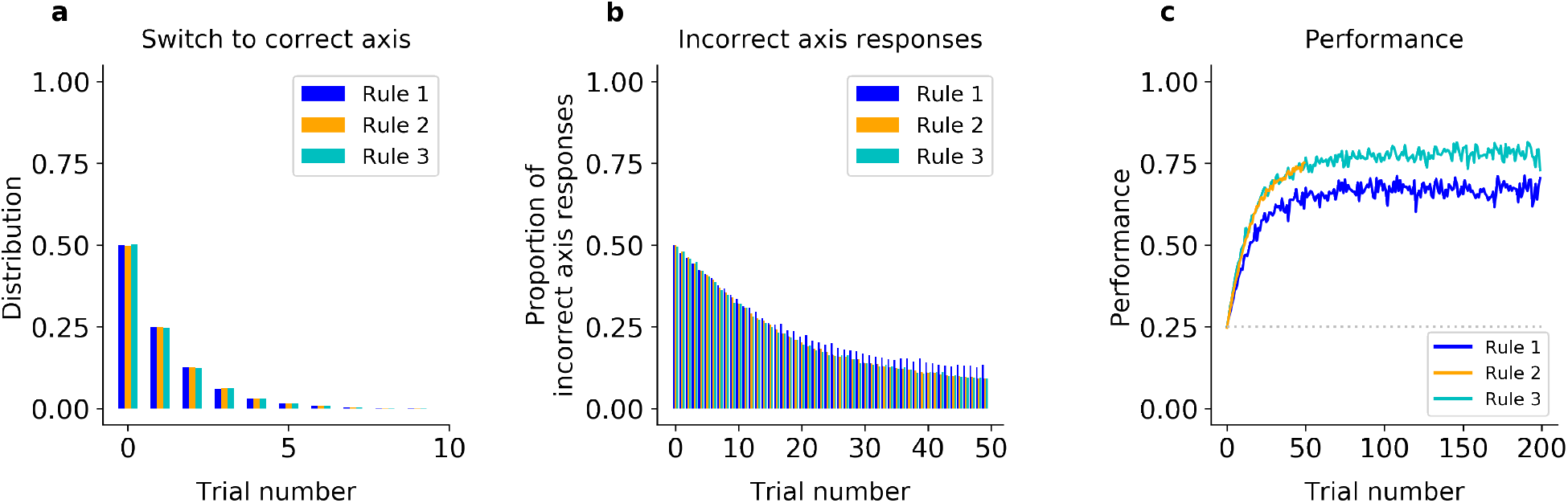
Incremental learner (QL) model fitted on Monkey S behavior (see Fig. S2 for Monkey C). (a) Trial number of the first response of the model on the correct axis after a block switch (compare to Fig. 1g, inset). (b) Proportion of responses of the model on the incorrect axis for the first 50 trials of each block (compare to Fig. 1g). (c) Model performance for each rule (averaged over blocks, compare to Fig. 1e).

In addition, because of the need to relearn feature-response associations after each block switch, the incremental QL model was unable to capture the dichotomy between the monkeys’ slower learning in Rule 1 and 3 (which share Axis 1) and the faster learning of Rule 2 (using Axis 2). As noted above, the monkeys performed Rule 2 at near asymptotic performance from the beginning of the block but were slower to learn which feature to attend to on blocks of Rule 1 and Rule 3 (Fig. 1g,h). In contrast, because the model learned each rule in the same way, the incremental learner performed similarly on all three rules (Fig. 2c for Monkey S, S2c for Monkey C). In particular, it performed correctly on the first trial in only 25% of Rule 2 blocks (Fisher test against behavior: p<10^(−4)^), and reached only 62% after 20 trials (p<10^(−4)^). As a result, on the first 20 trials, the difference in average percent performances was only Δ=4.0 between Rule 2 and Rule 1, and was only Δ=0.76 between Rule 2 and Rule 3 (similar results were seen when fitting the model to Monkey C, see Fig. S2). The same pattern of results was seen when the initial weights were free parameters (see Methods).

Altogether, these results argue simple incremental relearning of the axes and features *de novo* cannot reproduce the animals’ ability to instantaneously relearn the correct axis after a block switch or the observed differences in learning speed between the rules.

### Pure inference of previously learned rules cannot capture the behavior

The results above suggest that incremental learning is too slow to explain the quick switch between response axes displayed by the monkeys. So, we tested whether a model that leverages Bayesian inference can capture the animals’ behavior. A fully informed Bayesian ideal observer model (IO, see Fig. S1 and Methods) uses statistical inference to continually estimate which of the three rules is in effect, accumulating evidence (‘beliefs’) for each rule based on the history of previous stimuli, actions, and rewards. The IO model chooses the optimal action for any given stimulus, by averaging the associated actions’ values under each rule, weighted by the estimated likelihood that each rule is in effect. Like incremental learning, the IO model learns and changes behavior depending on experience. However, unlike incremental models, this model leverages perfect knowledge of the rules to learn rapidly, limited only by stochasticity in the evidence. Here, noisy stimulus perception is the source of such stochasticity, limiting both the speed of learning and asymptotic performance. Indeed, the IO model predicts the speed of initial (re)learning after a block switch should be coupled to the asymptotic level of performance. Furthermore, given that perceptual noise is shared across rules, the IO model also predicts the speed of learning will be the same for rules that use the same features.

As expected, when fit to the animal’s behavior, the IO model reproduced the animals’ ability to rapidly infer the correct axis (Fig. S3a,b,d,e). For example, when fitted to Monkey S behavior, the model initially responded on Axis 2 almost always immediately after each block switch cue (96% in all rules, Fisher test against monkey’s behavior p>0.4). Then, if this was incorrect, the model typically switched to the correct axis within 5 trials on 89% of blocks of Rule 1, and 95% of blocks of Rule 3 (Fisher test against monkey behavior: p>0.2 in both rules). The model maintained the correct axis with very few off-axis responses throughout the block (after trial 20, 1.3% in Rule 1; 1.1% in Rule 2; 1.2% in Rule 3; Fisher test against monkey’s behavior: p>0.6 in all rules).

However, the IO model could not capture the observed differences in learning speed for the different rules (Fig. S3c,f). To understand why, we looked at performance as a function of stimulus difficulty. As expected, the monkey’s performance depended on how difficult it was to categorize the stimulus (i.e., the morph level; psychometric curves shown in Fig. 3a-c for Monkey S, Fig. S4a-c for Monkey C). For example, in color blocks (Rule 2 and 3), the monkeys performed better for a ‘prototype’ red stimulus than for a ‘morphed’ orange stimulus (Fig. 3a-c). Indeed, on ‘early trials’ (first 50 trials) of Rule 2, Monkey S correctly responded to 96% (CI=[0.95,0.97]) of prototype stimuli, and only to 91%, CI=[0.90,0.92] of ‘morphed’ stimuli (p<10^(−4)^; similar results for Monkey C in Fig. S4). Rule 3 had a similar ordering: Monkey S correctly responded to 80% (CI=[0.77,0.82]) of prototype stimuli, and only 62%, CI=[0.60,0.64] of ‘morphed’ stimuli (p<10^(− 4)^). This trend continued as the animal learned Rule 3 (trials 50 to 200; 89%, CI=[0.88,0.90] and 74%, CI=[0.73,0.75], respectively for prototype and morphed stimuli, p<10^(−4)^).

**Figure 3:**
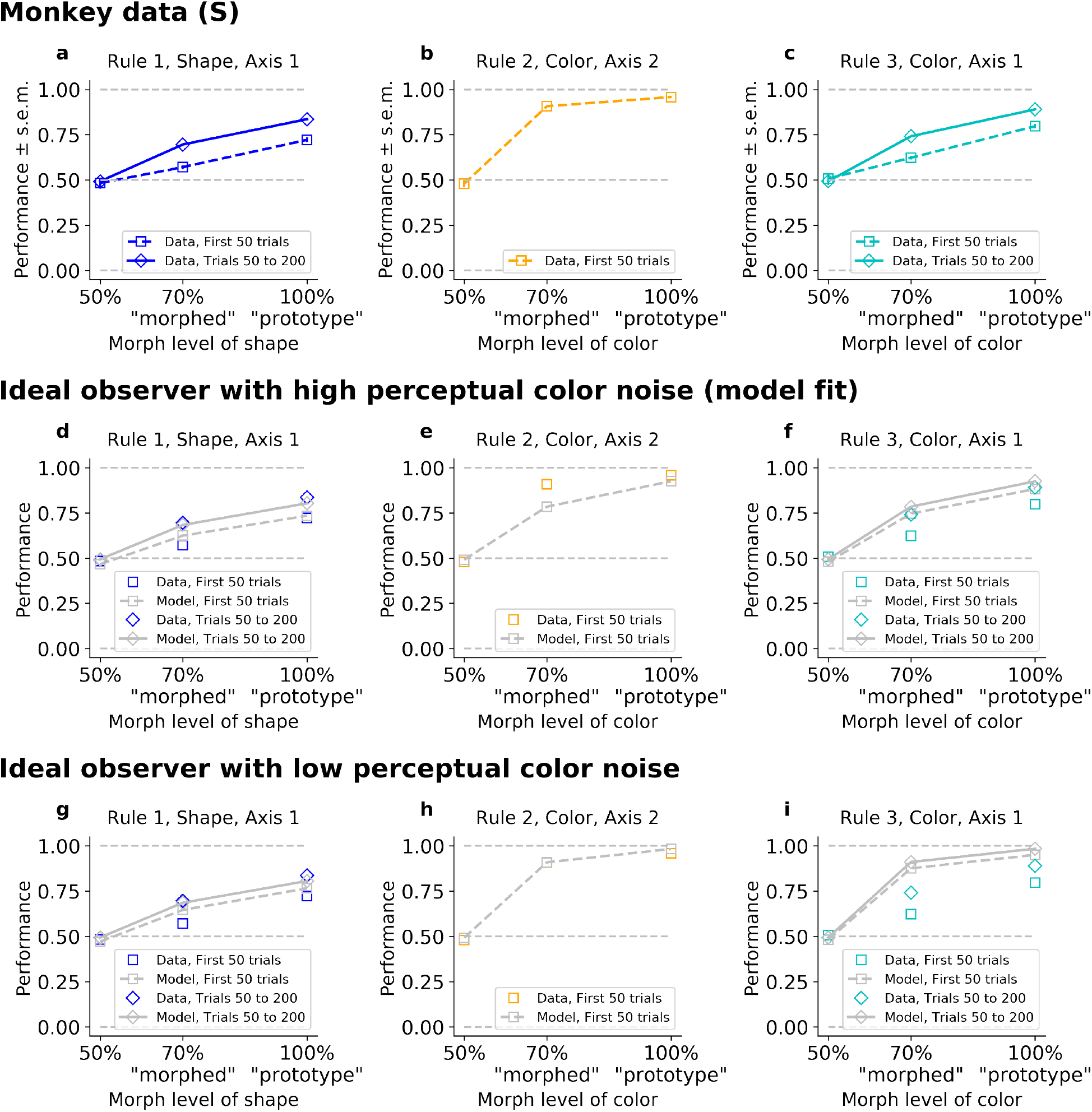
The ideal observer (IO), slow or fast, but not both. Fitted on Monkey S behavior (see Fig. S3 for Monkey C). Note that data were collapsed across 50%/150%; 30%/70%/130%/170%; and 0%/100% (non-collapsed psychometric functions can be seen in Fig. 5). (a-b-c) Performance for Rule 1, 2, and 3, as a function of the morphed version of the relevant feature. (d,e,f) Performance for Rule 1, 2, and 3, for IO model with high color noise. This parameter regime corresponds to the case where the model is fitted to the monkey’s behavior (see Methods). (j,k,l) Performance for Rule 1, 2, and 3, for IO model with low color noise. Here, we fixed κ_C_=6.

Importantly, there was a discrepancy between the performance on ‘morphed’ stimuli in Rule 2 versus Rule 3, with a difference in average percent performance of Δ=28 for the first 50 trials in both rules (p<10^(−4)^). This was still true, even if we considered Rule 2 against the last trials of Rule 3 (Δ=17, p<10^(−4)^). The same discrepancy was observed between the performance on ‘prototype’ stimuli in Rule 2 versus Rule 3, with a difference in average percent performance on Δ=16 for the first 50 trials in both rules (p<10^(−4)^), and Δ=6.8 if comparing to the last trials of Rule 3 (p<10^(−4)^).

The IO model captured the performance ordering on morphed and prototype stimuli for each rule (Fig. 3d-i, similar results for the model reproducing Monkey C, Fig. S4). However, the model performed similarly for morphed stimuli on Rule 2 and Rule 3. This is because both rules involve categorizing color and so they shared the same perceptual noise, leading to the same likelihood of errors. Furthermore, because perceptual noise limits learning and asymptotic performance in the IO model, it predicts the speed of learning should be shared across Rule 2 and Rule 3, and initial learning in both rules on the first 50 trials should be coupled to the asymptotic performance. Given this, the model had to trade-off between behavioral performance in Rule 2 and Rule 3. Using best- fit parameters, the model reproduced the animals’ lower asymptotic performance in Rule 3 by increasing color noise, and so it failed to capture the high performance on Rule 2 early on (Fig. 3e,f, Fig. S4e,f). The resulting difference in average percent performance for ‘morphed’ stimuli was only Δ=4.0 for the first 50 trials and Δ=0.0044 if we considered the last trials of Rule 3 (respectively Δ=4.2 and Δ=0.032 for ‘prototype’). Conversely, if we forced the model to improve color perception (by reducing perceptual noise, Fig. 3h,i, and Fig S4h,i), then it was able to account for the monkeys’ performance on Rule 2, but failed to match the animals’ behavior on Rule 3. The resulting difference in average percent performance was again only Δ=3.4 for the first 50 trials, and Δ=-0.17 if we considered the last trials of Rule 3 (respectively Δ=3.2 and Δ=-0.16 for ‘prototype’).

One might be concerned that including a correct generative prior on the transition between axes given by the specific task structure would solve this issue, as a Rule 2 block is always following a Rule 1 or Rule 3 block, hence possibly creating an inherent discrepancy in learning Rule 2 versus Rule 1 and Rule 3. However, the limiting factor was not the speed for axis discovery (which was nearly instantaneous, cf above), but the shared perceptual color noise between Rule 2 and Rule 3, coupling initial learning to asymptotic performance. Such a modified ideal observer could not account for the behavior (Fig. S5).

### The key features of monkeys’ behavior are reproduced by a hybrid model composing inference over axes and incremental relearning over features

To summarize, the main characteristics of the animals’ behavior were 1) rapid learning of the axis of response after a block switch, 2) immediately high behavioral performance of Rule 2, the only rule on Axis 2, and 3) slower relearning of Rules 1 and 3, which mapped different features onto Axis 1. Altogether, these results suggest that the animals learned axes and features separately, with fast learning of the axes and slower learning of the features. One way to conceive this is as a Bayesian inference model (similar to IO), but relaxing the assumption that the animal had perfect knowledge of the underlying rules (i.e., all of the stimulus-action-reward contingencies). We propose that the animals maintained two latent states (e.g., one corresponding to each axis of response) instead of the three rules we designed. Assuming each state had its own stimulus-action-reward mappings, the mappings would be stable for Rule 2 (Axis 2) but continually re-estimated for Rules 1 and 3 (Axis 1). To test this hypothesis, we implemented a hybrid model that inferred the axis of response while incrementally learning which features to attend for that response axis (Hybrid Q Learner, ‘HQL’ in the Methods, Fig. S1). In the model, the current axis of response was inferred through Bayesian evidence accumulation (as in the IO model), while feature-response weights were incrementally learned for each axis of response.

Intuitively, this model could explain all three core behavioral observations. First, inference allows for rapid switching between axes. Second, because only Rule 2 mapped to Axis 2, the weights for Axis 2 did not change and so the model was able to perform well on Rule 2 immediately. Third, because Rules 1 and 3 shared an axis of response, and, thus, a single set of feature-response association weights, this necessitated relearning associations for each block, reflected in the animal’s slower learning for these rules.

Consistent with this intuition, the HQL model provided an accurate account of the animals’ behavior. First, unlike the QL model, the HQL model reproduced the fast switch to the correct axis (Fig. 4a,b and Fig. S6a,b and Fig. S7a-c,g-i). Fitted to Monkey S behavior, the model initially responded on Axis 2 immediately after each block switch cue (91% in Rule 1, 89% in Rule 2 and Rule 3, Fisher test against monkey’s behavior p>0.05). Then, if this was incorrect, the model switched to the correct axis within 5 trials on 91% of blocks of Rule 1 and Rule 3 (Fisher test against behavior: p>0.2 in both rules). Similar to the animals, the model maintained the correct axis with very few off-axis responses throughout the block (on trial 20, 1.4% in Rule 1; 1.3%, in Rule 2; 1.5% in Rule 3, Fisher test against monkey’s behavior: p>0.7 in all rules).

**Figure 4:**
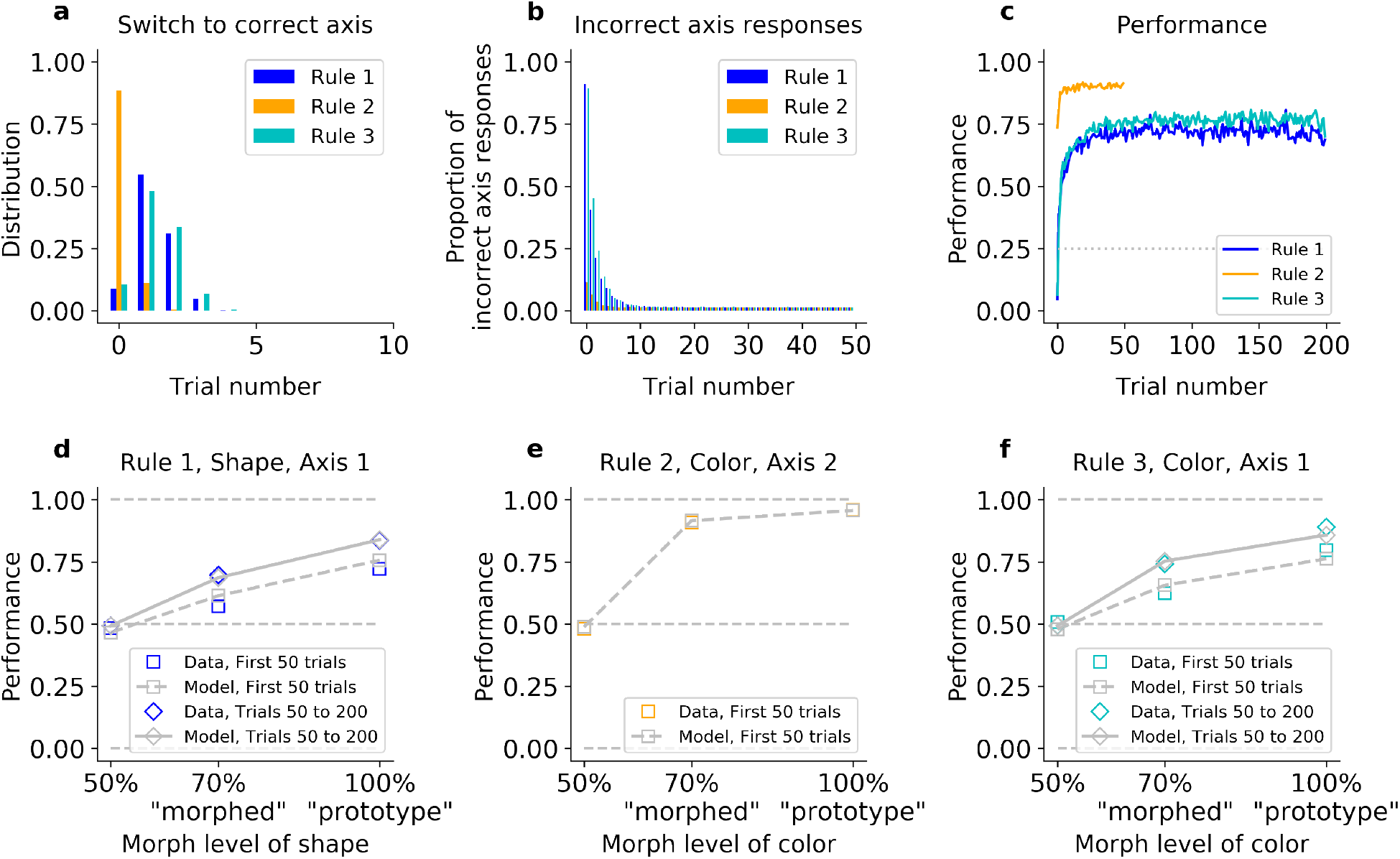
The hybrid learner (HQL) accounts both for fast switching to the correct axis, and slow relearning of Rule 1 and Rule 3. Model fit on Monkey S, see Fig. S6 for Monkey C. (a) Trial number for the first response on the correct axis after a block switch, for the model (compare to Fig. 1e inset). (b) Proportion of responses on the incorrect axis for the first 50 trials of each block, for the model (compare to Fig. 1e). (c) Performance of the model for the three rules (compare to Fig. 1g). (d,e,f) Performance for Rule 1, Rule 2 and Rule 3, as a function of the morphed version of the relevant feature.

Second, contrary to the IO model, the HQL model captured the animals’ fast performance on Rule 2 and slower performance on Rules 1 and 3 (Fig. 4c and Fig. S6c). As detailed above, animals were significantly better on Rule 2 than Rules 1 and 3 on the first 20 trials. The model captured this difference: fitted on Monkey S’s behavior, the difference in average percent performance on the first 20 trials was Δ=31 between Rule 2 and Rule 1, and Δ=29 between Rule 2 and Rule 3 (a Fisher test against monkey’s behavior gave p>0.05 for the first trial, p>0.1 for trial 20).

Third, the HQL model captured the trade-off between the animals’ initial learning rate and asymptotic behavioral performance in Rule 2 and Rule 3 (Fig. 4d-f and S6d-f). Similar to the animals, the resulting difference in average percent performance for ‘morphed’ stimuli was Δ=26 for the first 50 trials (Δ=16 if we considered the last trials of Rule 3; Δ=19 and Δ=9.9 for early and late ‘prototype’ stimuli, respectively). The model was able to match the animals’ performance because the weights for Axis 2 did not change from one Rule 2 block to another (Fig. S7e,k), and the estimated perceptual noise of color was low in order to account for the high performance of both morphed and prototype stimuli (Fig. 4e and Fig. S6e). To account for the slow re-learning observed for Rules 1 and 3, the best-fitting learning rate for feature-response associations was relatively low (Fig. 4d,f and Fig. S6d,f, Fig. S7d,f,j,l, Table S1).

### The effect of stimulus congruency (and incongruency) provides further evidence for the hybrid model

To further understand how the HQL model outperforms the QL and IO models, we examined the animal’s behavioral performance as a function of the relevant and irrelevant stimulus features. The orthogonal nature of the features and rules meant that stimuli could fall into two general groups: congruent stimuli had features that required the same response for both Rule 1 and Rule 3 (e.g., a green bunny, Fig. 1) while incongruent stimuli had features that required opposite responses between the two rules (e.g., a red bunny). Consistent with previous work^59–63^, the animals performed better on congruent stimuli than incongruent stimuli (Fig. 5a for Monkey S, Fig. S8a for Monkey C). This effect was strongest during learning, but persisted throughout the block (Fig. S9a,e): during early trials of Rules 1 and 3, the monkeys’ performance was significantly higher for congruent stimuli than for incongruent stimuli (gray vs. red squares in Fig. 5b; 94%, CI=[0.93,0.95] versus 57%, CI=[0.55,0.58] respectively; Δ=37, Fisher test p<10^(−4)^; see Fig. S8b for Monkey C). Similarly, the animals were slower to respond to incongruent stimuli (Fig. S10, Δ=25ms in reaction time for incongruent and congruent stimuli, t-test, p<10^(−4)^). In contrast, the congruency of stimuli had no effect during Rule 2 – behavior depended only on the stimulus color, suggesting the monkeys ignored the shape of the stimulus during Rule 2, even when the morph level of the color was more difficult (gray vs. red squares in Fig 5c; performance was 92%, CI=[0.90,0.93], and 93%, CI=[0.92,0.93] for congruent and incongruent stimuli, respectively; with Δ=-0.73; Fisher test, p=0.40; see Fig. S8c for Monkey C).

**Figure 5:**
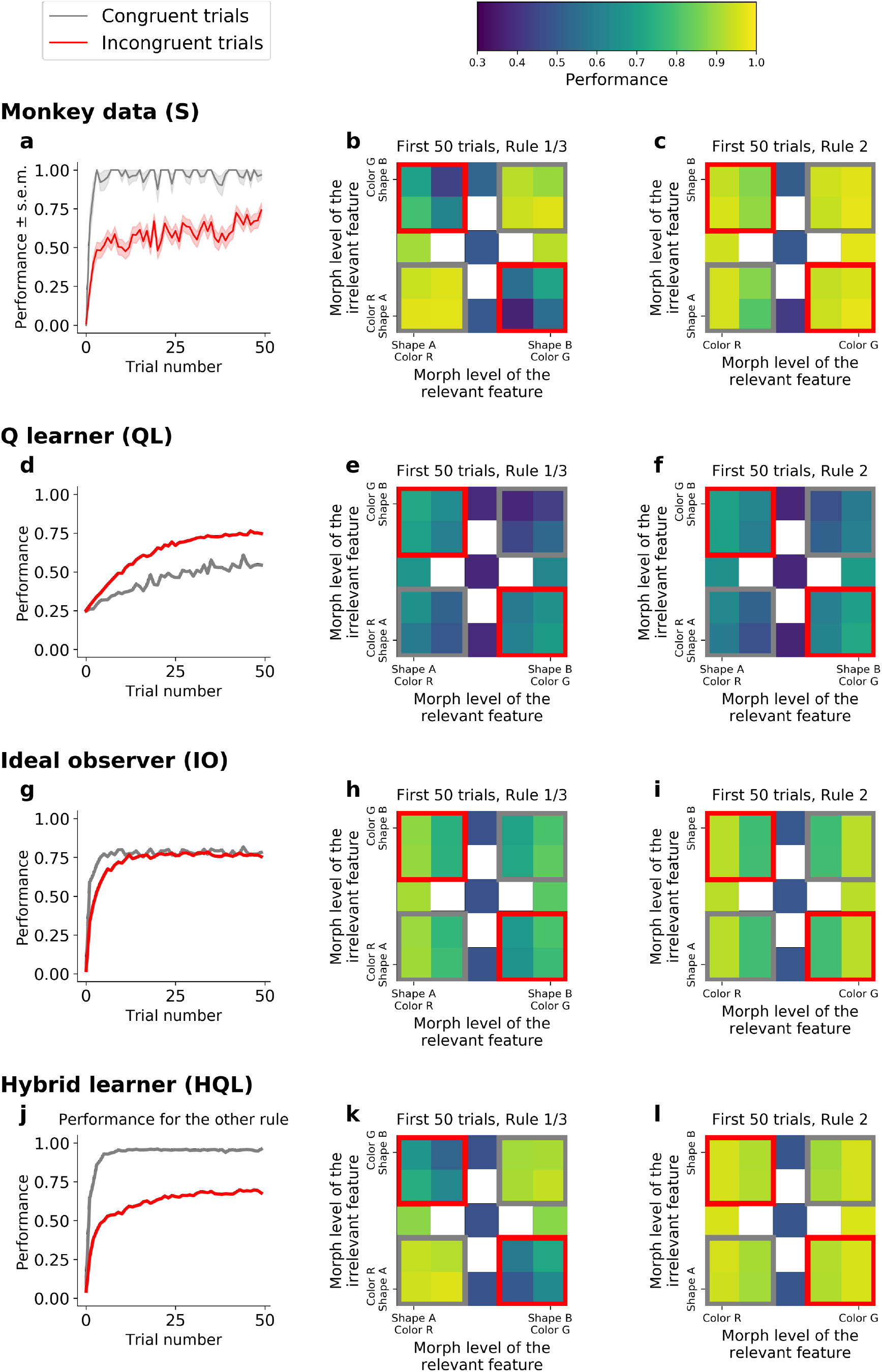
Comparison of incongruency effects in Monkey S and behavioral models (QL, IO, and HQL models). (a) Performance as a function of trial number for Rule 1 and Rule 3 (combined), for congruent and incongruent trials. (b) Performance for Rule 1 and 3 (combined, first 50 trials), as a function of the morph level for both color (relevant) and shape (irrelevant) features. Gray boxes highlight congruent stimuli, red boxes highlight incongruent stimuli. (c) Performance for Rule 2, as a function of the morph level for both color (relevant) and shape (irrelevant) features. Note the lack of an incongruency effect. (d,e,f) Same as a-c but for the QL model. (g,h,i) Same as a-c for the IO model. (j,k,l) Same as a-c but for the HQL model.

This incongruency effect provided further evidence for the HQL model. First, pure incremental learning by the QL model did not capture this result, but instead predicted an opposite effect. This is because incongruent trials were four times more likely than congruent trials (see Methods). As the QL model encodes the statistics of the task through error-driven updating of action values, the proportion of congruent vs. incongruent trials led to an anti-incongruency effect – the QL model fit to Monkey S predicted worse performance on congruent than incongruent trials (Fig. 5d,e; 45% and 62%, respectively; Δ=-16; Fisher test p<10^(−4)^; see Fig. S8d,e for Monkey C). Furthermore, for the same reason, the QL model produced a difference in performance during Rule 2 (Fig. 5f; 54% for congruent versus 64% for incongruent; Δ=-10; Fisher test p<10^(−4)^; see Fig. S8f for Monkey C, see also Fig. S9b,f for this effect throughout the block).

Second, the IO model also did not capture the incongruency effect. In principle, incongruency effects can be seen in this type of model when perceptual noise is large, because incongruent stimuli are more ambiguous when the correct rule is not yet known. However, given the statistics of the task, learning in the IO model quickly reached asymptotic performance, for both congruent and incongruent trials (Fig. 5g,h; 75% and 72% respectively; Δ=3.8 only; Fig. S8g,h for Monkey C), hence not reproducing the incongruency effect.

In contrast to the QL and IO models, the hybrid HQL model captured the incongruency effect. In the HQL model, the weights for mapping congruent stimuli to responses were the same for Rules 1 and 3. In contrast, the weights for incongruent stimuli must change for Rules 1 and 3. Therefore, the animals’ performance was immediately high on congruent stimuli, while the associations for incongruent stimuli had to be relearned on each block (Fig. S11). The model fitted to Monkey S behavior reproduced the greater performance on congruent than incongruent stimuli (Fig. 5g,h; 92% and 61%, respectively, Δ=31; see Fig. S8g,h for Monkey C). As with the monkey’s behavior, this effect persisted throughout the block (Fig. S9d,h). Finally, the HQL model captured the absence of incongruency effect in Rule 2 (Fig. 5i, green versus red squares; 92% and 93%, respectively; Δ=-1.0; see Fig. S8i for Monkey C), as there was no competing rule, there was no need to update the Axes 2 weights between blocks. As a result, only a hybrid model performing both rule switching of axis and rule-learning of features could account for the incongruency effect observed in the behavior.

## Discussion

In the present study, we investigated rule learning in two monkeys trained to switch between three category-response rules. Critically, the animals were not instructed as to which rule was in effect (only that the rule had changed). We compared two classes of models that were able to perform the task: incremental learning and inferential rule switching. Our results suggested that neither model fit the animals’ performance well. Incremental learning was too slow to capture the monkeys’ rapid learning of the response axis after a block switch. It was also unable to explain the high behavioral performance on Rule 2, which was the only rule requiring responses along the second axis. On the other hand, inferential learning was unable to reproduce the difference in performance for the two rules that required attending to the same feature of the stimulus (color), but responding on different axes (Rule 2 and Rule 3). Finally, when considered separately, neither of these two classes of models could explain the monkeys’ difficulty in responding to stimuli that had incongruent responses on the same axes (Rule 1 and Rule 3). Instead, we found that a hybrid model that inferred axes quickly and relearned features slowly, was able to capture the monkeys’ behavior. This suggests the animals were learning the current axis of response using fast inference while continuously re-estimating the stimulus-response mappings within an axis.

We assessed the generative performance of the hybrid model and falsified^64^ incremental learning and inferential rule switching considered separately. The superior explanatory power of the hybrid model suggests that the animals performed both rule switching and rule learning – even in a well- trained regime in which they could, in principle, have discovered perfect rule knowledge. The model suggests that the monkeys discovered only two latent states (corresponding to the two axes of response) instead of the three rules we designed, forcing them to perpetually relearn Rules 1 and 3. These two latent states effectively encode Rule 2 (alone on its response axis) on one hand, and a combination of Rule 1 and Rule 3 (sharing a response axis) on the other hand. The combination of rules in the second latent state caused the monkeys to continuously update their knowledge of the rules’ contingencies (mapping different stimulus features to actions). At first glance, one might be concerned that our major empirical finding about the discrepancy between fast switching and the slow updating of rules was inherent to the task structure. Indeed, a block switch predictably corresponds to a switch of axis, but does not always switch the relevant stimulus feature. Moreover, the features were themselves morphed, creating ambiguity when trying to categorize them and use past rewards for feature discovery. We can thus expect inherently the slower learning of the relevant feature. However, our experiments with ideal observer models demonstrate that an inference process reflecting the different noise properties of axes and features cannot by itself explain the two timescales of learning. The key insight is that, if slow learning about features is simply driven by their inherent noisiness, then the asymptotic performance level with these features should reflect the same degree of ambiguity. However, these do not match. This indicates that the observed fast and slow learning was not a mere representation of the generative model of the task.

The particular noise properties of the task may, however, shed light on a related question raised by our account: why the animals failed to discover the correct three-rule structure, which would clearly support better performance in Rule 1 and 3 blocks. Their failure to do so could shed light on the brain’s mechanisms for representing task properties and for discovering, splitting, or differentiating different latent states on the basis of their differing stimulus-action-response contingencies. One possible explanation is that the overlap between Rules 1 and 3 (sharing an axis of response) makes them harder to differentiate than Rule 2. In particular, the axis is the most discriminatory feature (being discrete and also under the monkey’s own explicit control), whereas the stimulus-reward mappings are noisier and continuous. Alternatively, a two-latent state regime may be rational given the cognitive demands of this task, including the cost of control or constraints on working memory when routinely switching between latent states (stability-flexibility trade-off)^63,65,66^. Although more difficult, perhaps training the monkeys on the four possible rules permissible by the experimental design (and thus adding Rule 4, sharing the axis of Rule 2 but using the feature shape, cf. missing rule in Fig. 1d) would have forced encoding latent states from implicit information, including stimuli features, not restricted by the dimensionality of motor responses. Finally, the monkeys may have encoded each rule as a separate latent state with a different training protocol (e.g., longer training, a higher ratio of incongruent versus congruent trials, or less morphed and more prototyped stimuli).

Understanding how the brain discovers and manipulates latent states would give insight into how the brain avoids catastrophic interference. In artificial neural networks, sequentially learning tasks causes catastrophic interference, such that the new task interferes (overwrites) the representation of the previously learned task^67^. In our task, animals partially avoided catastrophic interference by creating two latent states where learning was independent. For example, learning stimulus-response mappings for Rules 1 and 3 did not interfere with the representation of Rule 2. In contrast, Rules 1 and 3 did interfere – behavior was re-learned on each block. Several solutions to this problem have been proposed in machine learning literature such as orthogonal subspaces^68^ and generative replay^69^. Similarly, recent advances in deep reinforcement learning have started to elucidate the importance of incorporating metalearning in order to speed up learning and to avoid catastrophic interference^70,71^. Our results suggest the brain might solve this problem by creating separate latent states where learning is possible within each latent state. How these latent states are instantiated in the brain is an open question, and discovering those computations promises exciting new insights for algorithms of learning.

Finally, our characterization of the computational contributions of rule switching and rule learning, and the fortuitous ability to observe both interacting in a single task, leads to a number of testable predictions about their neural interactions. First, our results make the strong prediction that there should be two latent states represented in the brain – the representation for the two rules competing on one axis (Rules 1 and 3) should be more similar to one another than to the neural representation of the rule alone on the other axis (Rule 2). This would not be the case if the neural activity was instead representing three latent causes. Furthermore, our hybrid model suggests there may be a functional dissociation for rule switching and rule learning, such that they are represented in distinct networks. One hypothesis is that this dissociation is between cortical and subcortical regions. Prefrontal cortex may carry information about the animal’s trial beliefs (i.e. over the two latent states) in a similar manner as perceptual decision making when accumulating evidence from noisy stimuli^72–75^. Basal ganglia may, in turn, be engaged in the learning of rule-specific associations^1–10^. Alternatively, despite their functional dissociation, future work may find both rule switching and rule learning are represented in the same brain regions (e.g., prefrontal cortex).

## Acknowledgments

The authors thank Sam Zorowitz for helpful discussions on the statistical modeling platform Stan. This research was supported by U.S. Army Research Office ARO W911NF-16-1-047 (ND) and NIMH R01MH129492 (TJB).

## Author contributions

Conceptualization: F.B., S.T., M.M., T.B., N.D.

Methodology: F.B., S.T., M.M., T.B., N.D.

Software: F.B.

Validation: F.B., S.T., M.M., T.B., N.D.

Formal analysis: F.B.

Investigation: F.B., S.T.

Resources: T.B.

Data curation: S.T., T.B.

Writing – original draft preparation: F.B.

Writing – review and editing: F.B., S.T., M.M., T.B., N.D.

Visualization: F.B.

Supervision: T.B., N.D.

Project administration: T.B., N.D.

Funding acquisition: T.B., N.D.

## Declaration of interests

The authors declare no competing interests.

## Data and materials availability

Codes supporting the findings of this study will be available on GitHub (https://github.com/buschman-lab/InterplayRuleLearningRuleSwitching) and Code Ocean. All data supporting the findings of this paper will be available on Dryad.

## Materials and Methods

### 1 Experimental design and model notation

Two adult (8-11 years old) male rhesus macaques (Macaca mulatta) participated in a category-response task. Monkeys S and C weighed 12.7kg and 10.7kg respectively. All experimental procedures were approved by Princeton University Institutional Animal Care and Use Committee and were in accordance with the policies and procedures of the National Institutes of Health.

Stimuli were rendering of three dimensional models that were built using POV-Ray and MATLAB (Mathworks). They were presented on a Dell U2413 LCD monitor positioned at a viewing distance of 58cm. Each stimulus was generated with a morph-level drawn from a circular continuum (Eq. 1) between two prototype colors *C* (red and green) and two prototype shapes *S* (“bunny” and “tee”; Fig. 1b).

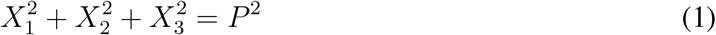

Where X is the parameter value in a feature dimension e.g. L,a,b values in CIELAB color space. Radius(P) was chosen such that there was enough visual discriminability between morph-levels. Morph-levels in shape dimension were built by circular interpolation of the parameters defining the lobes of the first prototype with the parameters defining the corresponding lobes of the second prototype. Morph-levels in the color dimension were built by selecting points along photometrically isoluminant circle in CIELAB color space that connected red and green prototype colors. We used percentage to quantify the deviation of each morph-level from prototypes (0% and 100%) on the circular space. Morph-levels between 0% and 100% correspond to – *π* to 0, and morph-levels 100% to 200% correspond to 0 to *π* on the circular space. Morph-levels for color and shape dimension were generated at 8 levels: 0%, 30%, 50%, 70%, 100%, 130%, 150%, 170%. 50% morph-levels for one feature (color or shape) were only generated for prototypes for the other feature (shape or color respectively). The total stimulus set consisted of 48 images. By creating a continuum of morphed stimuli, we could independently manipulate stimulus difficulty along each dimension.

Monkeys were trained to perform three different rules 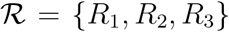 (Fig.1 c,d). All of the rules had the same general structure: the monkeys categorized a visual stimulus according to its shape or color, and then responded with a saccade, *α* ∈ *Actions*, to one of four locations *Actions* = {*Upper* – *Left, Upper* – *Right, Lower* – *Left, Lower* – *Right*} (Fig. 1a). Each rule required the monkeys to attend-to and categorize either the color or shape feature of the stimulus, and then respond with a saccade along an axis 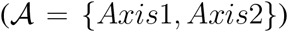. *Axis 1* corresponded to Upper-Left, Lower-Right locations and Axis 2 corresponded to Upper-Right, Lower-Left locations. As such, the correct response depended on the rule in effect and stimulus presented during the trial. Rule 1 (*R*_1_) required a response on Axis 1, to the Upper-Left or Lower-Right locations when the stimulus was categorized as “bunny” or “tee”, respectively. Rule 2 (*R*_2_) required a response on Axis 2, to the Upper-Right or Lower-Left locations when the stimulus was categorized as red or green, respectively. Rule 3 (*R*_3_) required a response on Axis 1, to the Upper-Left or Lower-Right locations when the stimulus was categorized as green or red, respectively. In this way, the three rules were compositional: Rule 2 and Rule 3 shared the response to the same feature of the stimulus (color) but different axes. Similarly, Rule 1 and Rule 3 shared the response to same axis (*Axis 1*), but to different features.

The monkeys initiated each trials by fixating a dot on the center of the screen. During a fixation period (lasting 500-700 ms), the monkeys were required to maintain their gaze within a circle with radius of 3.25 degrees of visual angle around the fixation dot. After the fixation period, the stimulus and all four response locations were displayed simultaneously. The monkeys made their response by breaking fixation and saccading to one of the four response locations. Each response location was 6 degrees of visual angle from the fixation point, located at 45°, 135°, 225°, and 315° degrees relative to vertical. The stimulus diameter was 2.5 degrees of visual angle. The animal’s reaction time was taken as the moment of leaving the fixation window, relative to the onset of the stimulus. Trials with a reaction time lower than 150ms were aborted, and the monkey received a brief timeout. Following a correct response, monkeys were provided with a small reward, while incorrect responses led to a brief timeout (*r* ∈ {0,1}). Following all trials, there was an intertrial interval of 2 to 2.5 seconds before the next trial began. The time distributions were adjusted according to task demands and previous literature^76^.

Note that both Rule 1 and Rule 3 required a response along the same axis (*Axis 1*). Half of the stimuli were “congruent”, such that they led to the same response for both rules (e.g., a green bunny is associated with an Upper-Left response for both rules). The other half of stimuli were “incongruent”, such that they led to different responses for both rules (e.g., a red bunny is associated with a Upper-Left and Lower-Right response, for Rule 1 and Rule 3, respectively). To ensure the animals were performing the rule, incongruent stimuli were presented on 80% of the trials.

Animals followed a single rule for a “block” of trials. After the animal’s behavioral performance on that rule reached a threshold, the task would switch to a new block with a different rule. The switch between blocks was triggered when the monkeys’ performance was greater than or equal to 70% on the last 102 trials of Rule 1 and Rule 3 or the last 51 trials of Rule 2. For Monkey S, each performance, for “morphed” and “prototype” stimuli independently, had to be above threshold. Monkey C’s performance was weaker, and a block switch occurred when the average performance for all stimuli was above threshold. Also, to avoid that Monkey C perseverated on one rule for an extended period of time on a subset of days, the threshold was reduced to 65% over the last 75 trials for Rule 1 and Rule 3, after 200 or 300 trials. Switches between blocks of trials were cued by a flashing screen, a few drops of juice, and a long time out (50secs). Importantly, the rule switch cue did not indicate the identity of the rule in effect or the upcoming rule. Therefore, the animal still had to infer the current rule based on its history.

Given the limited number of trials performed each day and to simplify the task structure for the monkeys, the axis of response always changed following a block switch. During Axis *1* blocks, whether Rule 1 or Rule 3 was in effect was pseudo-randomly selected. These blocks were interleaved by Axis *2* blocks, which were always Rule 2. Pseudo-random selection of Rule 1 and Rule 3 within Axis 1 blocks was done to ensure the animal performed at least one block of each rule during each session (accomplished by never allowing for three consecutive blocks of the same rule).

As expected, given their behavioral performance, the average block length varied across monkeys: 50-300 trials for Monkey S, and 50-435 trials for Monkey C. Rule 1 and Rule 3 blocks were on average 199 trials for Monkey S and 222 trials for Monkey C. Rule 2 blocks were shorter because they were performed more frequently and were easier given the task structure. They were on average 56 trials for Monkey S and 52 trials for Monkey C. Overall, the behavioral data includes 20 days of behavior from Monkey S, with an average of 14 blocks per day, and 15 days for Monkey C, with an average of 6.5 blocks per day.

#### Additional details on training

Given the complex structure of the task, monkeys were trained for months until they fully learned the structure of the task and they could consistently perform at least 5 blocks each day. Monkeys learned the structure of the task in multiple steps. They were first trained to hold fixation and to associate stimuli with reward by making saccades to target locations. To begin with shape categorization (Rule 1), monkeys learned to associate monochrome versions of prototype stimuli with two response locations on *Axis 1.* Stimuli were then gradually colored by using an adaptive staircase procedure. To begin training on color categorization (Rule 2), monkeys learned to associate red and green squares with two response locations on Axis 2. Prototype stimuli gradually appeared on the square and finally replaced the square using an adaptive staircase procedure. After this stage, monkeys were trained to generalize across morph-levels in 5% morph-level steps using an adaptive staircase method until they could generalize up to 20% morph-level away from the prototypes, for color and shape features. Rule 3 was added at this stage with a cue (purple screen background). Once the monkey was able to switch between Rule 1 and Rule 3, the cue was gradually faded and finally removed. After monkeys learned to switch between three rules, the morph-levels 30% and 40% were introduced. Monkey S and monkey C were trained for 36 and 60 months, respectively. Behavioral data reported here are part of data acquisition during electrophysiological recording sessions. From this point, only behavioral sessions in which monkeys performed at least 5 blocks were included for further analysis. In order to encourage generalization for shape and color features, during non-recording days, monkeys were trained on the larger number of morph-levels (0%, 20%, 30%, 40%, 50%, 60%, 70%, 80%, 100%, 120%, 130%, 140%, 150%, 160%, 170%, 180%).

### 2 Modeling noisy perception of color and shape

All the models studied below model stimulus perception in the same way (Fig. S1), via a signal-detection-theory like account by which objective stimulus features are modeled as magnitudes corrupted by continuously distributed perceptual noise. In particular, the color and shape of each stimulus presented to the animals are either the prototype features *s_Tc_* ∈ {*red, green*} and *s_Ts_* ∈ {*bunny, tee*}, or a morphed version of them. The feature continuous spaces are projected onto the unit circle from –*π* to *π*, such that each stimulus feature has a unique radius angle, with prototype angles being 0 and *π*. The presented stimulus is denoted **s_M_** = (*s_Mc_, s_Ms_*). We hypothesize that the monkeys perceive a noisy version of it, denoted **s_K_** = (*s_Kc_, s_Ks_*). We model it by drawing two independent samples, one from each of two Von Mises distributions, centered around each feature (*s_Mc_* and *s_Ms_*, for color and shape), and parameterized by the concentrations *κ_c_* and *κ_s_* respectively. The models estimate each initial feature presented by computing its posterior distribution, given the perceived stimulus, i.e. by Von Mises distributions centered on *s_Kc_* and *s_Ks_*, with same concentrations *κ_c_* and *κ_s_*.

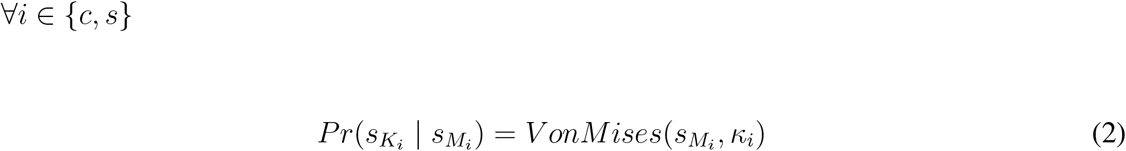

and so:

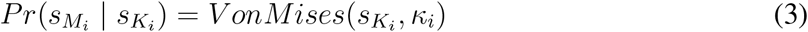

with:

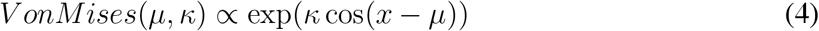

### 3 Modeling action selection

All the models studied below use the same action-selection stage. Given the perceived stimulus at each trial **s_K_** = (*s_Kc_*, an action is chosen so as to maximize the expected reward 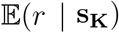 by computing max_*a*_ *Pr*(*r* = 1 | **s_K_**,*a*) which corresponds to maximizing the probability of getting a reward, given the perceived stimulus. We use the notation 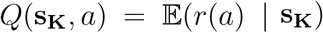 as in [54,77] and refer to these values as *Q values.* Note that even a deterministic “max” choice rule at this stage does not, in practice, imply noiseless choices, since *Q* depends on **s_K_**, and the perceptual noise in this quantity gives rise to variability in the maximizing action that is graded in action value, analogous to a softmax rule. For that reason, we do not include a separate choice-stage softmax noise parameter (which would be unidentifiable relative to perceptual noise *κ*), though we do nevertheless approximate the max choice rule with a softmax (but using a fixed temperature parameter), for implementational purposes (specifically, to make the choice model differentiable). To control the asymptotic error rate, we also include an additional probability of lapse (equivalently, adding “epsilon-greedy” choice). Altogether, two fixed parameters implement an epsilon-greedy softmax action-selection rule: the lapse rate *ϵ* and the inverse temperature *β*.

The action-selection rule is:

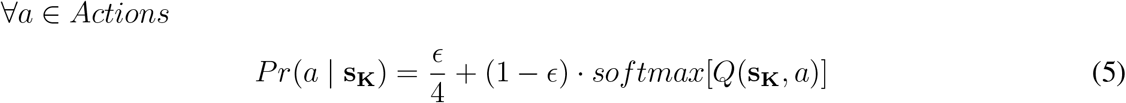

The lapse rate *ϵ* is directly estimated from the data, by computing the proportion of trials where the incorrect axis of response is chosen, asymptotically. It is evaluated to 0.02 for both Monkey S and Monkey C. The inverse temperature of the softmax *β* is also fixed (*β* = 10), to allow the algorithm to approximate the max while remaining differentiable (cf. use of Stan below).

### 4 Fit with Stan

All our models shared common noisy perceptual input and action selection stages (Fig. S1, and Methods). The models however differed in the intervening mechanism for dynamically mapping stimulus to action value (see Fig. S1). Because of noise perception at each trial (Eq. 2), and because the cumulative distribution function of a Von Mises is not analytic, the models are fitted with Monte Carlo Markov chains (MCMC) using Stan^78^. Each day of recording is fitted separately, and the mean and standard deviation reported in Table S1 are between days. Fitting scripts will be available on github upon acceptance. We validated convergence by monitoring the potential scale reduction factor R-hat (which was < 1.05 for all simulations) and an estimate of the effective sample size (all effective sample sizes > 100) of the models’ fits^79^.

Models’ plots correspond to an average of 1000 simulations of each day of the dataset (with the same order of stimuli presentation). Statistics reported in the article were done with Fisher’s exact test (except a t-test for reaction times, Fig. S10).

### 5 Incremental learner: QL model

This model corresponds to Fig. 2 (Monkey S) and S2 (Monkey C). It also appears in Fig. 5, S8, S9, S11.

In this model the agent is relearning each rule after a block as a mapping between stimuli and actions, by computing a stimulus-action value function as a linear combination of binary feature-response functions *ϕ*(**s_K_**, *a*) with feature-response weights w. This implements incremental learning while allowing for some generalization across actions. The weights are updated by the delta rule (*Q* learning with linear function approximation, see [80], chapter 9). The weights are reset from one block to the other, and the initial values for each reset are set to zero. Fitting them does not change the results (see paragraph 5.4 below).

#### 5.1 Computation of the feature-response matrix

Given a morph perception at trial *t*, **s_K_** = (*s_Kc_, s_Ks_*), a feature-response matrix is defined as:

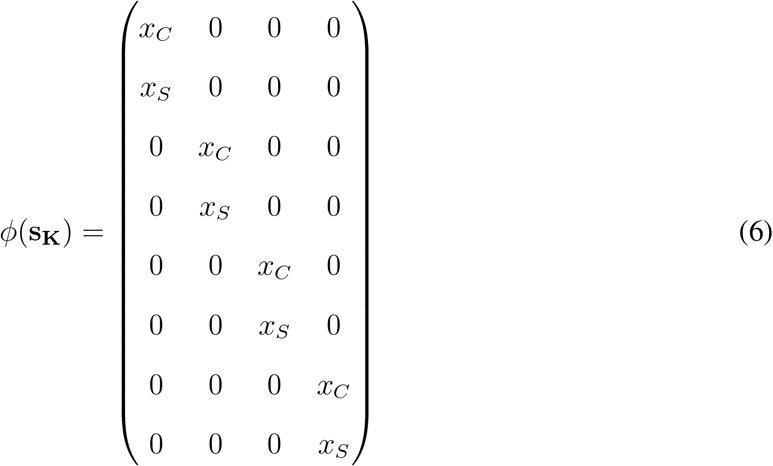

where *x_C_* ∈ {-1,1} depends on whether the perceived morph for color *s_Kc_* is classified as green or red (Eqs. 2 and 3), and *x_S_* ∈ {-1,1} whether the perceived morph for shape *s_Ks_* is classified as tee or bunny (Eqs. 2 and 3). In order for the algorithm to remain differentiable, we approximate {-1,1} with a sum of sigmoids. Each column of the matrix *ϕ*(**s_K_**) is written *ϕ*(**s_K_**, *a)* below and corresponds to an action *a* ∈ *Actions*.

#### 5.2 Linear computation of Q values and action selection

In order to compute Q values, the feature-response functions *ϕ*(**s_K_**, *a*) are weighted by the feature-response weight vector **w** = (*w*_1_,.., *w*_8_) (see [80], equation (9.8)):

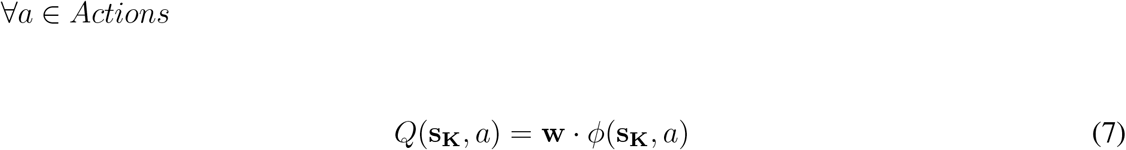

Action selection is done through the epsilon-greedy softmax rule (Eq. 5).

Thus asymptotic learning of Rule 1 would require **w** = [0,1, 0, −1, 0,0, 0, 0]. Learning Rule 2 would require **w** = [0, 0, 0, 0,−1,0,1, 0]. Learning Rule 3 would require **w** = [−1, 0,1, 0, 0, 0, 0, 0].

#### 5.3 Weight vector update

Once an action *a_t_* is chosen and a reward *r_t_* is received at trial *t,* the weights are updated by the delta rule with learning rate *α* (see [80], equation (9.7)).

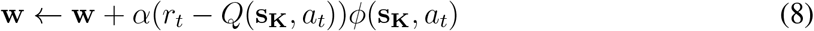

#### 5.4 Parameter values

*β* and e are fixed to respectively 10 and 0.02. See Table S1 for parameter values. As predicted from the behavior, noise perception is higher for shape than for color (*κ_C_* > *κ_S_*). The initial weight vector w_0_ is set to zero at the beginning of each block day. Fitting these weights instead gives the same results (as then **w**_0_ has a mean=[−0.070,0.031,0.093,−0.054,−0.081,−0.022,0.062,−0.0048] for Monkey **S** and **w**_0_ has a mean=[−0.099,0.070,0.069,−0.089,−0.053,−0.011,0.030,−0.0098] for Monkey C).

### 6 Optimal Bayesian inference over rules: IO model

This model corresponds to Fig. 3, and S3; also appears in Fig. 5, and S4, S5, S8, S9, S11.

In this model, we assume a perfect knowledge of combination mappings between prototype stimuli and actions as *rules.* Learning is discovering which rule is in effect by Bayesian inference. This is done through learning, over the trials, the probability for each rule to be in effect in a block (or *belief*) from the history of stimuli, actions and rewards. At each trial, this belief is linearly combined to the likelihood of a positive reward given the stimulus to compute a value for each action. This likelihood encapsulates knowledge of the three experimental rules. An action is chosen as per described above in Eq. 5. The beliefs over rules are then updated through Bayes rule using the likelihood of the reward received, given the chosen action and the stimulus perception. Once the rule is discovered, potential errors thus only depend on the possible miscategorization of the stimulus features (Eqs. 2 and 3), or eventually on exploration (Eq. 5).

#### 6.1 Belief over rules

The posterior probability of rule 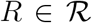 to be in effect in the block is called the belief over the rule *b*(*R*) = *Pr*(*R* | s_K_,*a, r*), given the perceived stimulus s_K_, the action *a* and the reward *r*. The beliefs **b**(*R*) at the beginning of each block are initialized to **b**_0_ = [*b*_1_, 1 – *b*_1_ – *b*_3_, *b*_3_] where *b*_1_ and *b*_3_ are fitted, to test for a systematic initial bias towards one rule.

#### 6.2 Computation of values

The beliefs are used to compute the **Q** values for the trial:

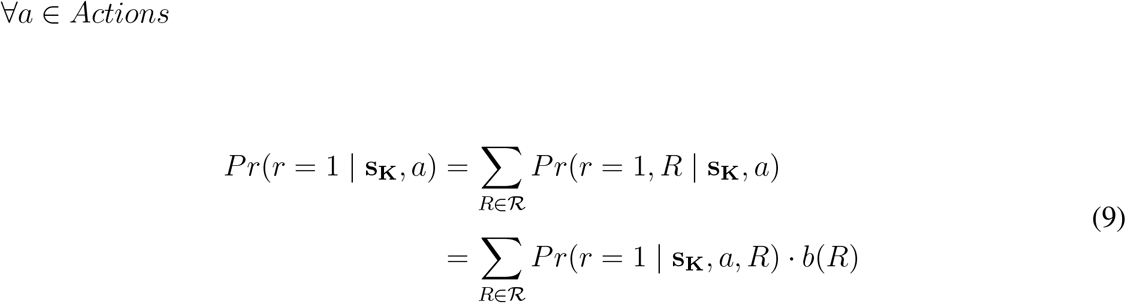

with (marginalization over the possible morph stimuli presented):

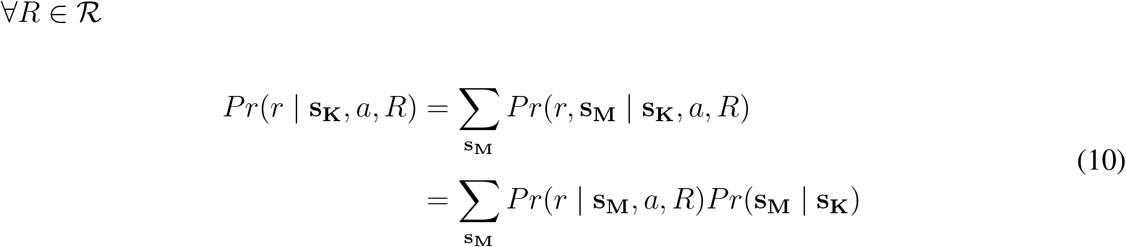

Noting *p_C_* = *p*(*s_Mc_* = *red*|*s_Kc_*); and *p_S_* = *p*(*s_Ms_* = *bunny* | *s_K_s*), gives:

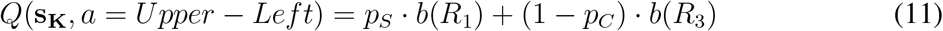

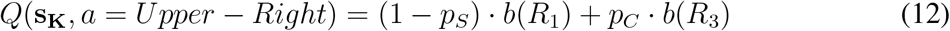

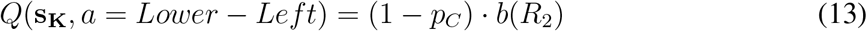

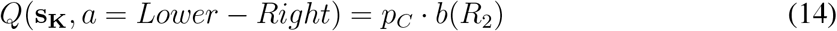

#### 6.3 Belief update

From making an action *a_t_* ∈ *Actions*, the agent receives a reward *r_t_* ∈ {0,1}, and the beliefs over rules are updated:

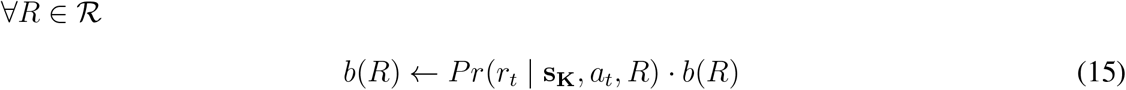

with *Pr*(*r_t_* |**s_K_**, *a_t_,R*) the likelihood of observing reward *r_t_* for the chosen action *a_t_*.

Note that because of the symmetry of the task, 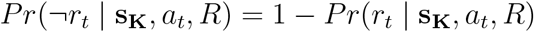.

#### 6.4 Parameter values

*β* and *ϵ* are respectively fixed to 10 and 0.02. See Table S1 for parameter values. As predicted from the behavior, there is an initial bias for Rule 2 for the model fitted on Monkey **S** behavior (*b*_2_ > *b*_3_ > *b*_1_). Also, noise perception is higher for shape than for color for both monkeys (*κ_C_* > *κ_S_*). In the version of the model with low perceptual color noise (Fig. 3 and S3), all the parameters remain the same, except that we fix *κ_C_* = 6 for all simulated days.

### 7 Hybrid incremental learner: HQL model

The hybrid incremental learner combines inference over axes with incremental learning, using a Q-learning with function approximation to relearn the likelihood of rewards given stimuli per axis of response. This model corresponds to Fig. 4 and S6; also appears in Fig. 5, S7, S8, S9, S11.

#### 7.1 Belief over axes

The posterior probability of an axis 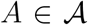 to be the correct axis of response in a block is called the belief over axis *b*(*A*) = *Pr*(*A* | **s_K_**, *a, r*), given the perceived stimulus **s_K_**, the action *a* and the reward *r*. The beliefs over axes are initialized at the beginning of each block to **b**_0_ = (*b_ax_*, 1 – *b_ax_*).

#### 7.2 Computation of the feature-response matrix

As for the incremental learner above, given a morph perception at trial *t*, **s_K_** = (*s_Kc_,s_Ks_*), a feature-response matrix is defined as:

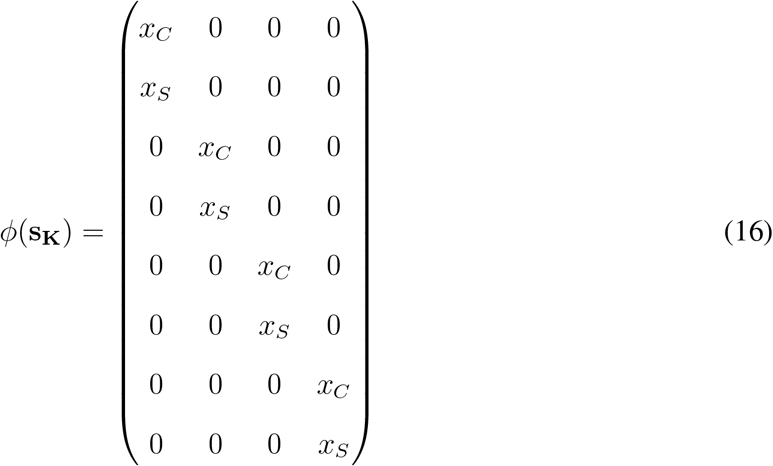

where *x_C_* ∈ {–1, 1} depends on whether the perceived morph for color *s_Kc_* is classified as green or red (Eqs. 2 and 3), and *x_s_* ∈ {–1,1} whether the perceived morph for shape *s_Ks_* is classified as tee or bunny (Eqs. 2 and 3). Each column of the matrix *ϕ*(**s_K_**) is written *ϕ*(**s_K_**, *a*) below and corresponds to an action *a* ∈ *Actions*.

#### 7.3 Computation of values

The beliefs are used to compute the Q values for the trial:

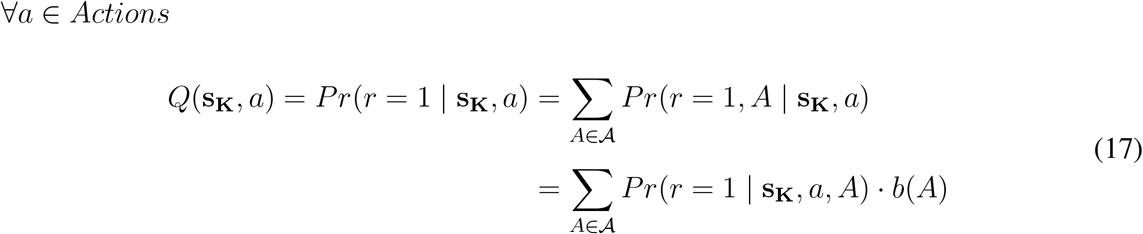

Contrary to the ideal observer, here the likelihood of reward per action *Pr*(*r* = 1 | **s_K_**, *a, A*) is learned through function approximation.

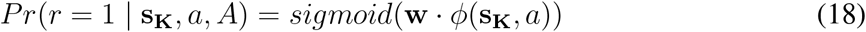

Action selection is done through the epsilon-greedy softmax rule (Eq. 5).

#### 7.4 Weight vector update

Once an action *a_t_* is chosen and a reward *r_t_* is received at trial *t*, the weights are updated through gradient descent with learning rate *a* (see [80], equation (9.7)).

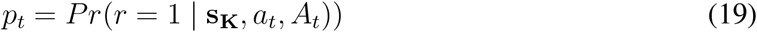

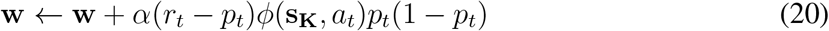

As learning improved steadily in this model contrary to the asymptotic behavior of monkeys, we implemented a weight decay to asymptotic values **w**_0_:

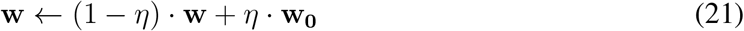

Note that resetting the weights at the beginning of each block and adding a weight decay (or a learning rate decay) provide similar fits to the dataset. Also, this decay can be included in the previous two models without any change of our results and conclusions.

#### 7.5 Belief update

From making an action *a_t_* ∈ *Actions*, the agent receives a reward *r_t_* ∈ {0,1}, and the beliefs over axes are updated:

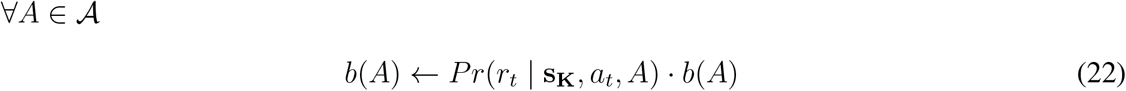

with *Pr*(*r_t_* | **s_K_**,*a_t_,A*) the likelihood of observing reward *r_t_* for the chosen action *a_t_*.

#### 7.6 Parameter values

*β* and *ϵ* are fixed to respectively 10 and 0.02. As predicted from the behavior, noise perception is higher for shape than for color for both monkeys (*κ_C_* > *κ_S_*). Also, the model fitted on Monkey S behavior has an initial bias for Axis2 (*b_ax_* < 0.5). For fitting the model on Monkey C behavior, we fix *b_ax_* = 0.5. Finally, the fitted values of **w**_0_ correspond to an encoding of an average between Rule 1 and Rule 3 on *Axis* 1, and an encoding of Rule 2 on *Axis*2, for both monkeys.

### 8 Research standards

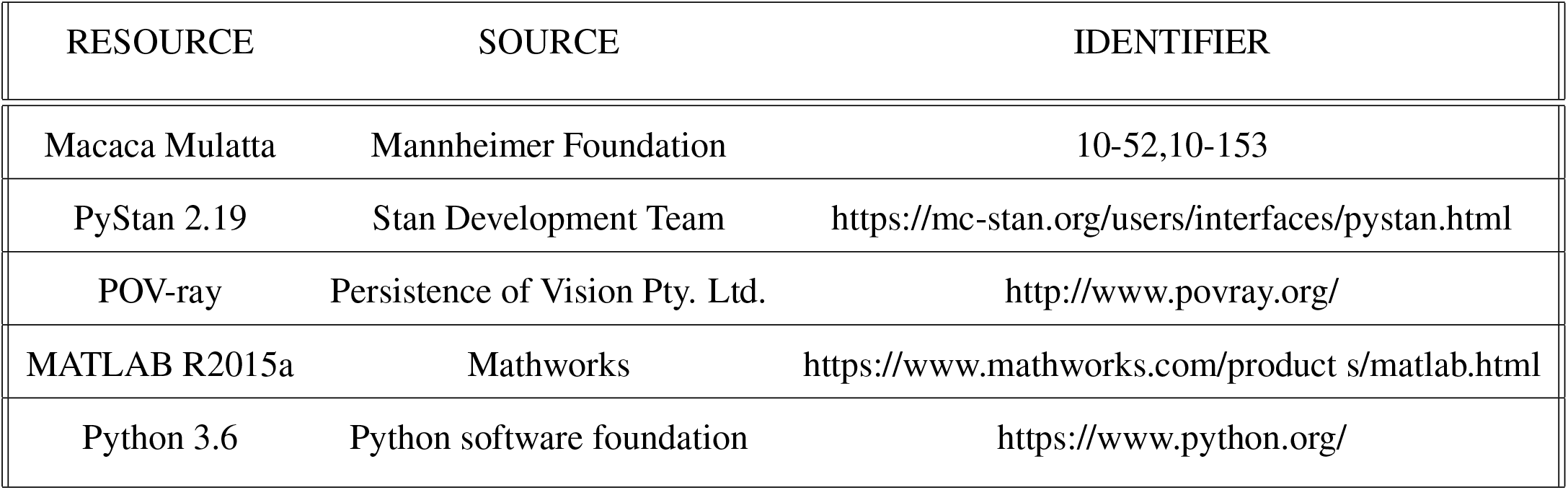

## Supplementary figures

**Figure S1:**
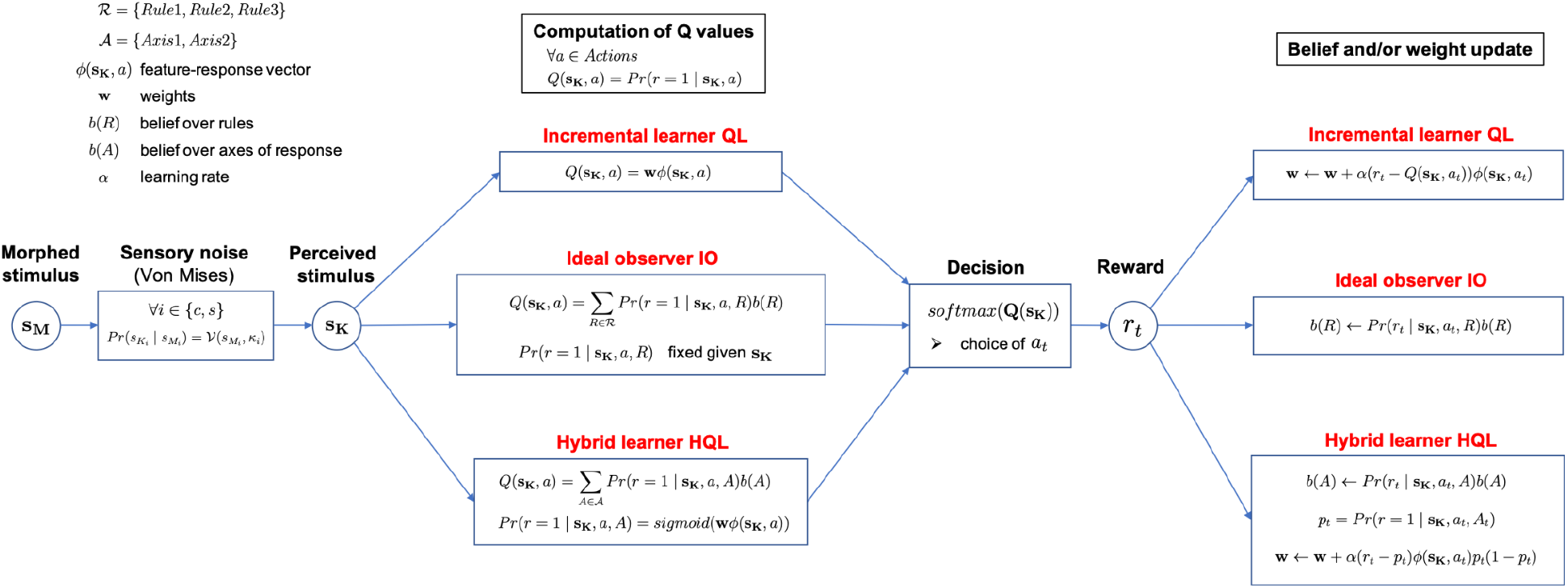
The 3 models.

**Figure S2:**
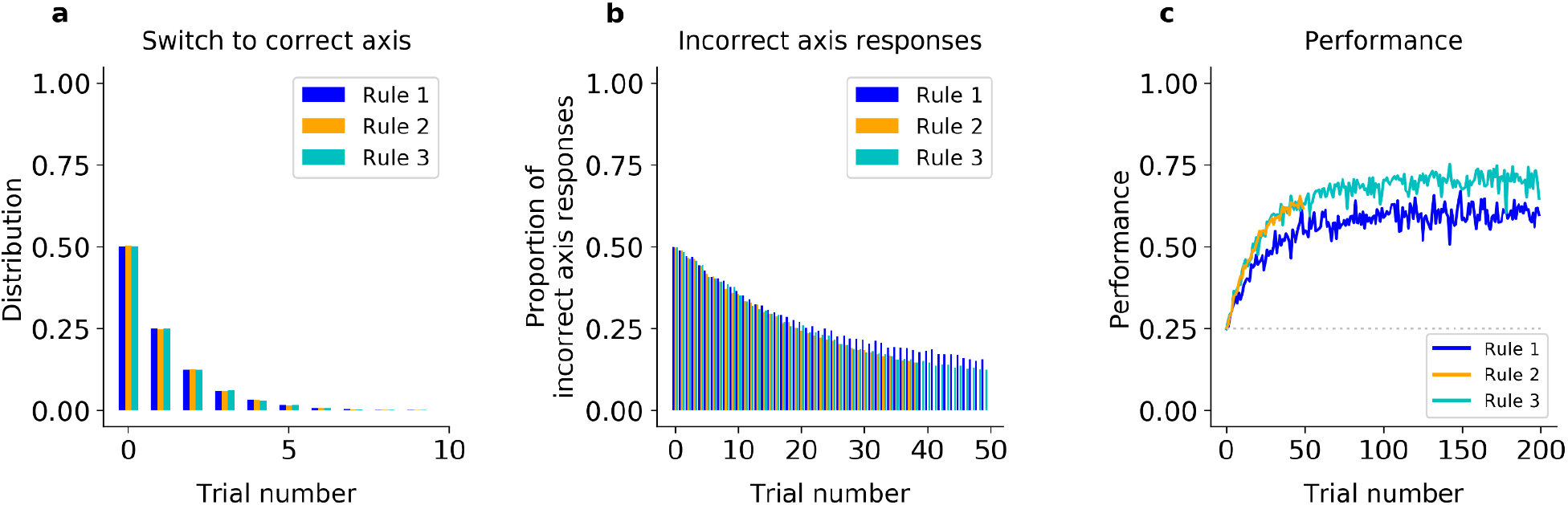
Incremental learner (QL) model fitted on Monkey C behavior (see Fig. 2 for Monkey S). (a) Trial number of the first response of the model on the correct axis after a block switch (compare to Fig. 1h, inset). (b) Proportion of responses of the model on the incorrect axis for the first 50 trials of each block (compare to Fig. 1h). (c) Model performance for each rule (averaged over blocks, compare to Fig. 1f). <u>Statistics of QL model fitted on Monkey C</u>: First, the model made a response on the correct axis on the first trial with a probability of only 50% in Rule 1, Rule 2 and Rule 3. The model performed 28% of off-axis responses after 20 trials in Rule 1, and 25% in Rule 2 and Rule 3. Second, the model performed correctly on the first trial in only 25% of Rule 2 blocks, and reached only 51% after 20 trials. As a result, on the first 20 trials, the difference in average percent performance was only Δ=3.3 between Rule 2 and Rule 1, and only Δ=-0.39 between Rule 2 and Rule 3.

**Figure S3:**
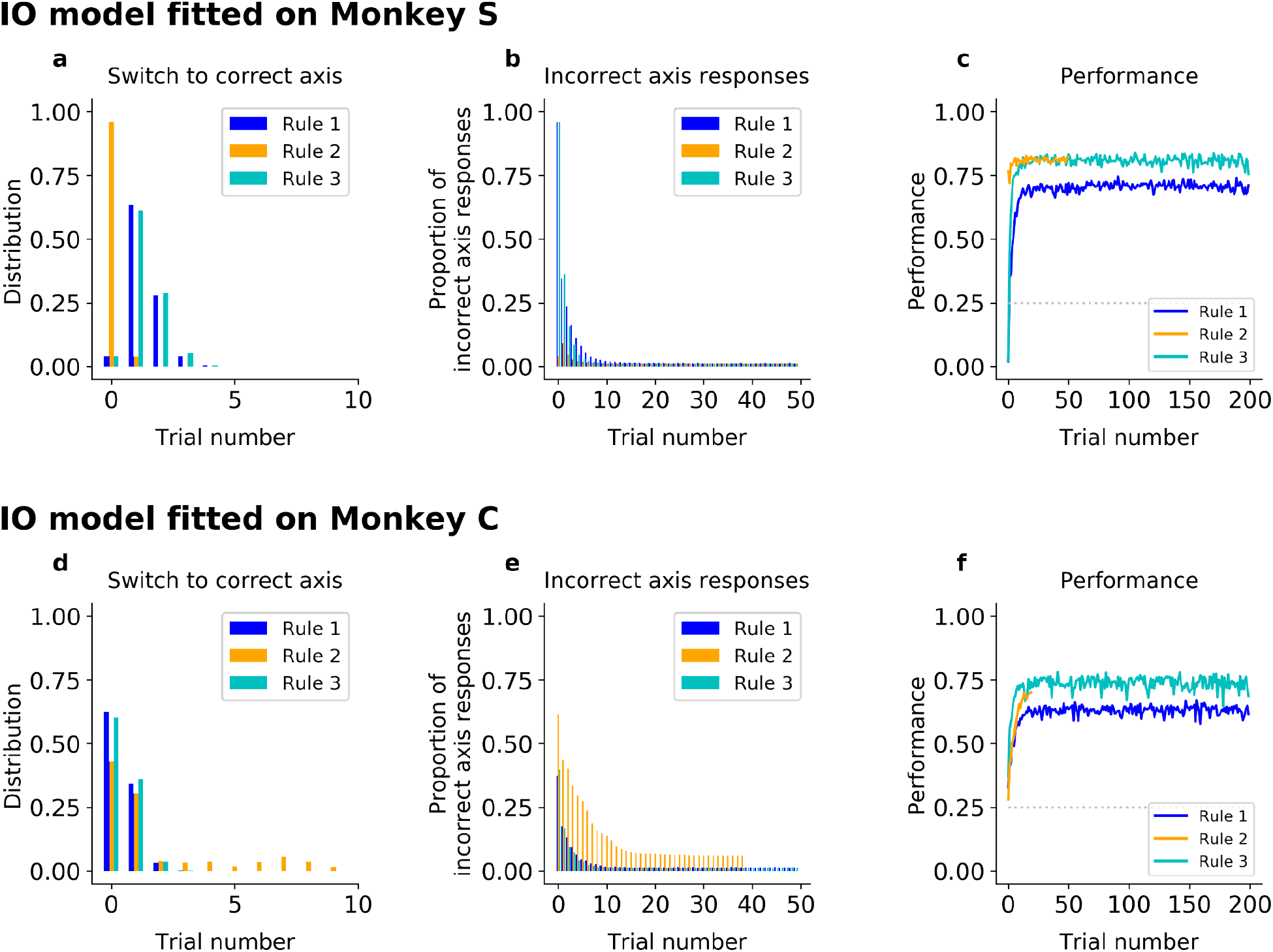
(a-c) IO model fitted on Monkey S behavior. (a) Trial number of the first response of the model on the correct axis after a block switch (compare to Fig. 1e, inset). (b) Proportion of responses of the model on the incorrect axis for the first 50 trials of each block (compare to Fig. 1e), (c) Model performance for each rule (averaged over blocks, compare to Fig. 1g). (d-f) Same, but for Monkey C. <u>Statistics of IO model fitted on Monkey C</u>: The model responded on the correct axis on the first trial with a probability of 63% in Rule 1, 38% in Rule 2, and 60% in Rule 3. The model maintained the correct axis with very few off-axis responses throughout the block (after trial 20, 1.3% in Rule 1; 6.8%, in Rule 2; 1.3% in Rule 3, Fisher test against monkey behavior: p>0.5 in all rules).

**Figure S4:**
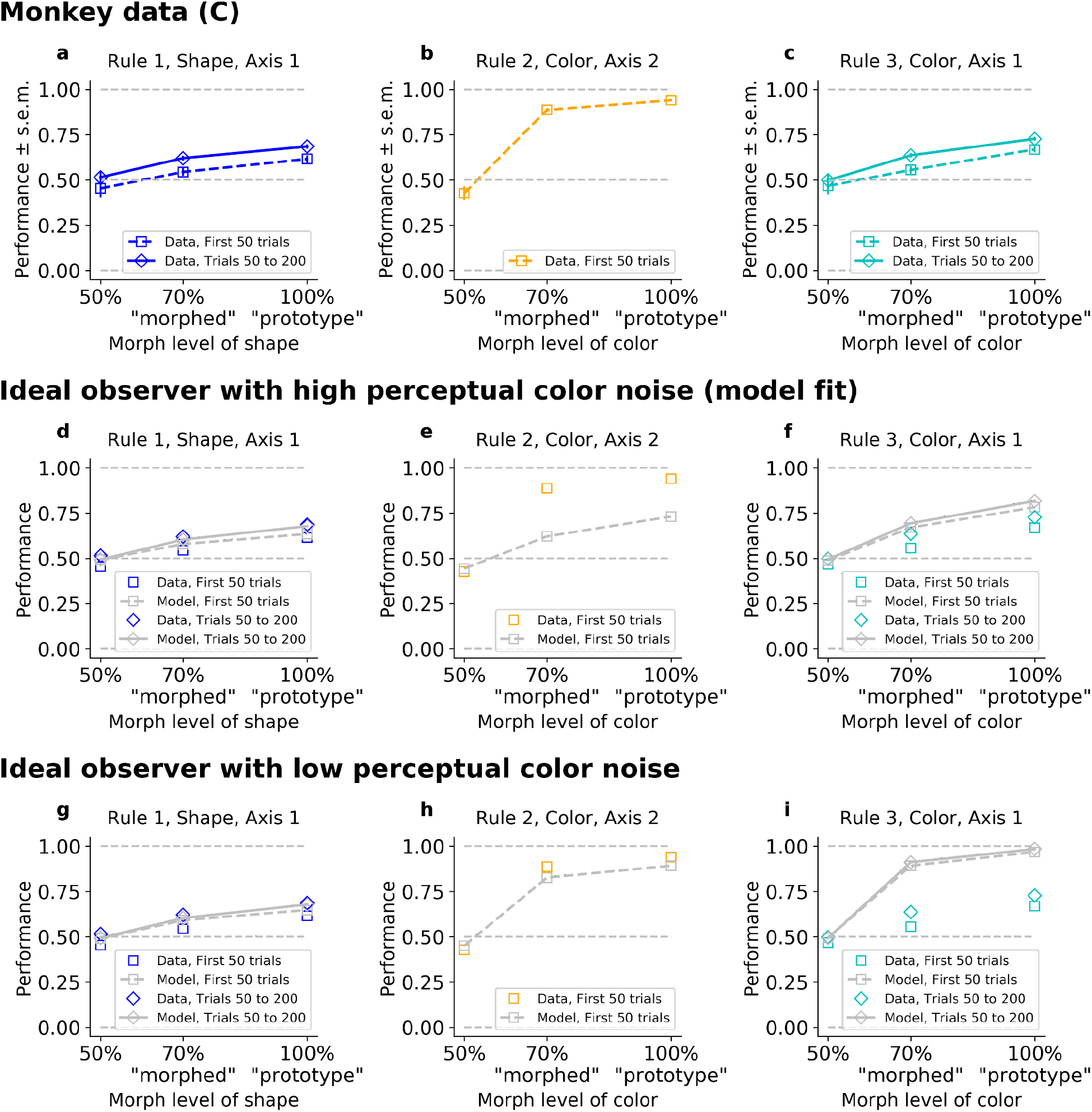
The ideal observer (IO), slow or fast, but not both. Fitted on Monkey C behavior (see Fig. 3 for Monkey S). (a-b-c) Performance for Rule 1, 2, and 3, as a function of the morphed version of the relevant feature. (d,e,f) Performance for Rule 1, 2, and 3, for IO model with high color noise. This parameter regime corresponds to the case where the model is fitted to the monkey’s behavior (see Methods). (j,k,l) Performance for Rule 1, 2, and 3, for IO model with low color noise. Here, we fixed KC=6. <u>Statistics on Monkey C</u>: There was a discrepancy between the performance of ‘morphed’ stimuli in Rule 2 versus Rule 3, with a difference in average percent performances of Δ=33 for the first 50 trials in both rules (p<10^-4^), and still Δ=24 if we considered Rule 2 against the last trials of Rule 3 (p<10^-4^). The same discrepancy was observed between the performance of ‘prototype’ stimuli in Rule 2 versus Rule 3, with a difference in average percent performances of Δ=27 for the first 50 trials in both rules (p<10^-4^), and still Δ=21 if we considered the last trials of Rule 3 (p<10^-4^). <u>Statistics of IO model fitted on Monkey C</u>: While the IO model, using best-fit parameters, reproduced poor asymptotic performance in Rule 3 by increasing color noise (low concentration), it then failed to capture the high performance on Rule 2 early on. The resulting difference in performance for ‘morphed’ stimuli was only Δ=4.8 for the first 50 trials and Δ=-7.2 if we considered the last trials of Rule 3 (respectively Δ=5.1 and Δ=-8.6 for ‘prototype’).

**Figure S5:**
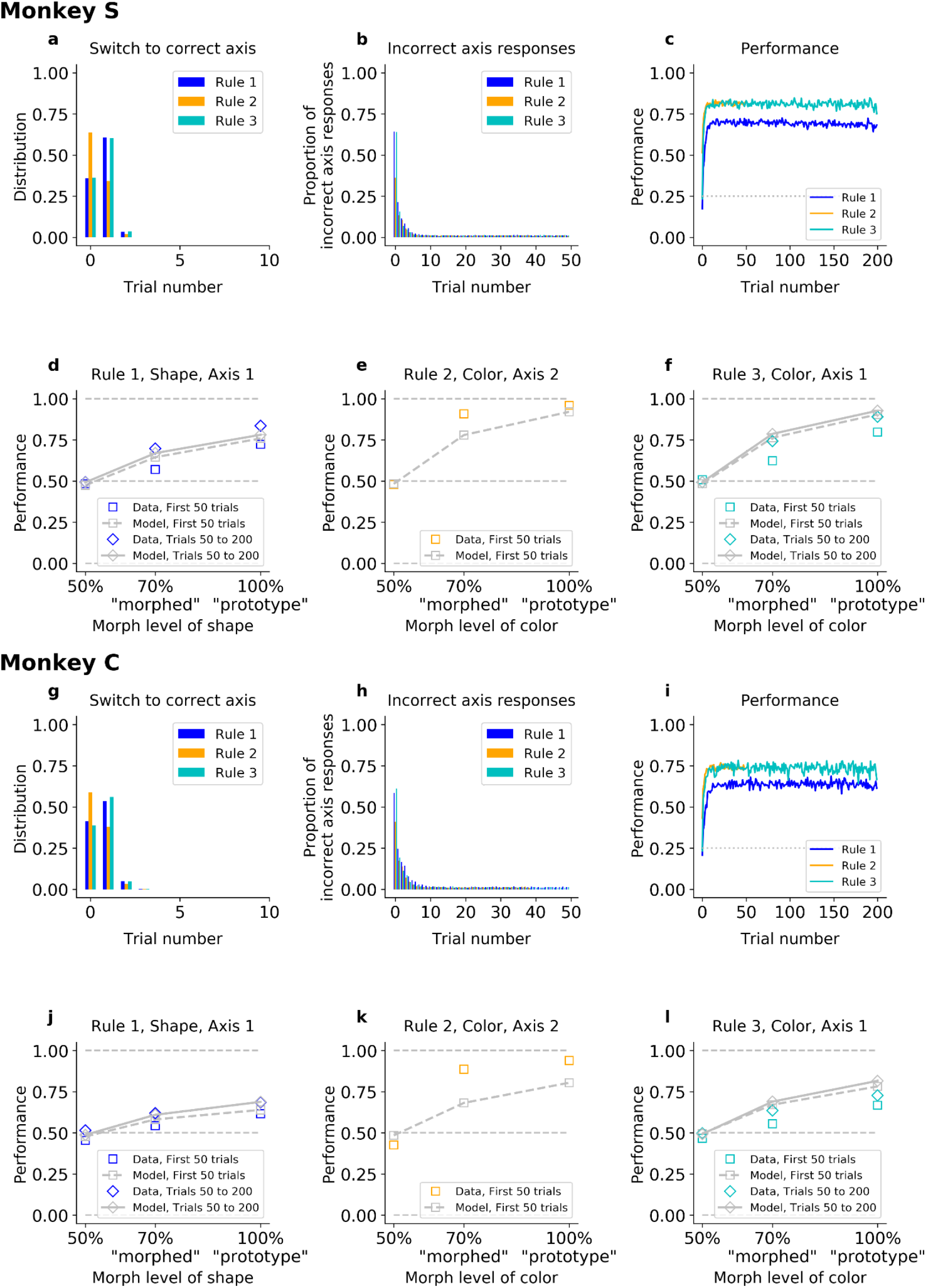
The ideal observer model including a correct generative prior on the transition between axes given by the specific task structure. This is defined by fixing the values of the initial belief states over rules to (b_1_=0; b_2_=1; b_3_=0) for a Rule 2 block, and (b_1_=0.5; b_2_=0; b_3_=0.5) for a Rule 1 or Rule 3 block. The model is fitted on Monkeys S and C.

**Figure S6:**
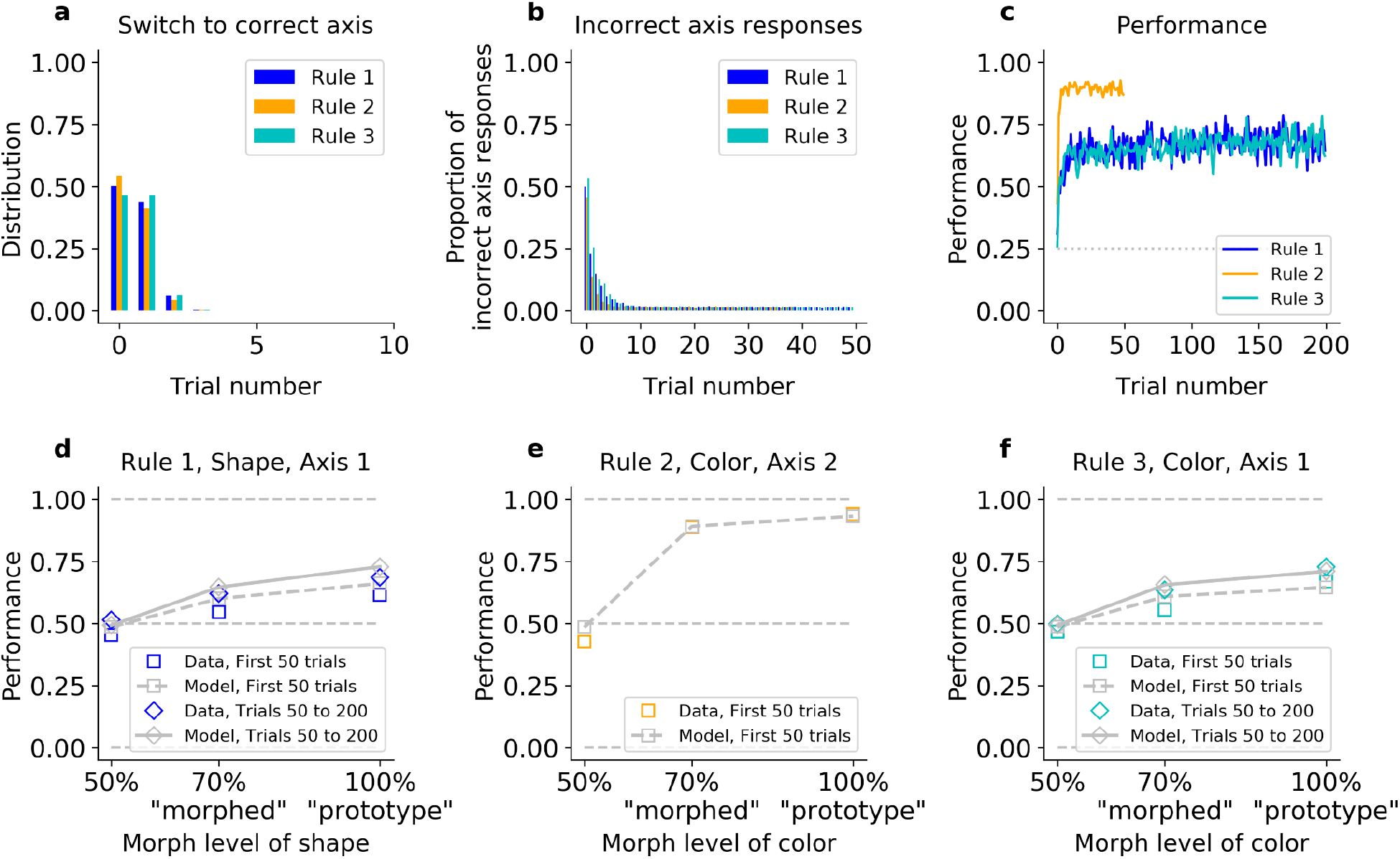
The hybrid learner (HQL) accounts both for fast switching to the correct axis, and slow relearning of Rule 1 and Rule 3. Model fit on Monkey C, see Fig. 4 for Monkey S. (a) Trial number for the first response on the correct axis after a block switch, for the model (compare to Fig. 1f inset). (b) Proportion of responses on the incorrect axis for the first 50 trials of each block, for the model (compare to Fig. 1f). (c) Performance of the model for the three rules (compare to Fig. 1h). (d,e,f) Performance for Rule 1, Rule 2 and Rule 3, as a function of the morphed version of the relevant feature. <u>Statistics of HQL model fitted on Monkey C</u>: First, The model responded on the correct axis on the first trial with a probability of 50% in Rule 1, 54% in Rule 2, and 47% in Rule 3. The model maintained the correct axis with very few off-axis responses throughout the block (after trial 20, 1.5% in Rule 1; 1.7%, in Rule 2; 1.5% in Rule 3, Fisher test against monkey’s behavior: p>0.5 in all rules). Second, the HQL model could capture the animal’s fast performance on Rule 2 and slower performance on Rules 1 and 3: the difference in average percent performances on the first 20 trials was Δ=28 both between Rule 2 and Rule 1 and between Rule 2 and Rule 3. Third, not only the HQL model captured the performance ordering on morphed and prototype stimuli for each rule separately, but the model was able to trade-off between initial and asymptotic behavioral performance in Rule 2 and Rule 3, for both ‘morphed’ and ‘prototype’ stimuli. The resulting difference in performance for ‘morphed’ stimuli was Δ=29 for the first 50 trials and Δ=24 if we considered the last trials of Rule 3 (respectively Δ=29 and Δ=22 for ‘prototype’).

**Figure S7:**
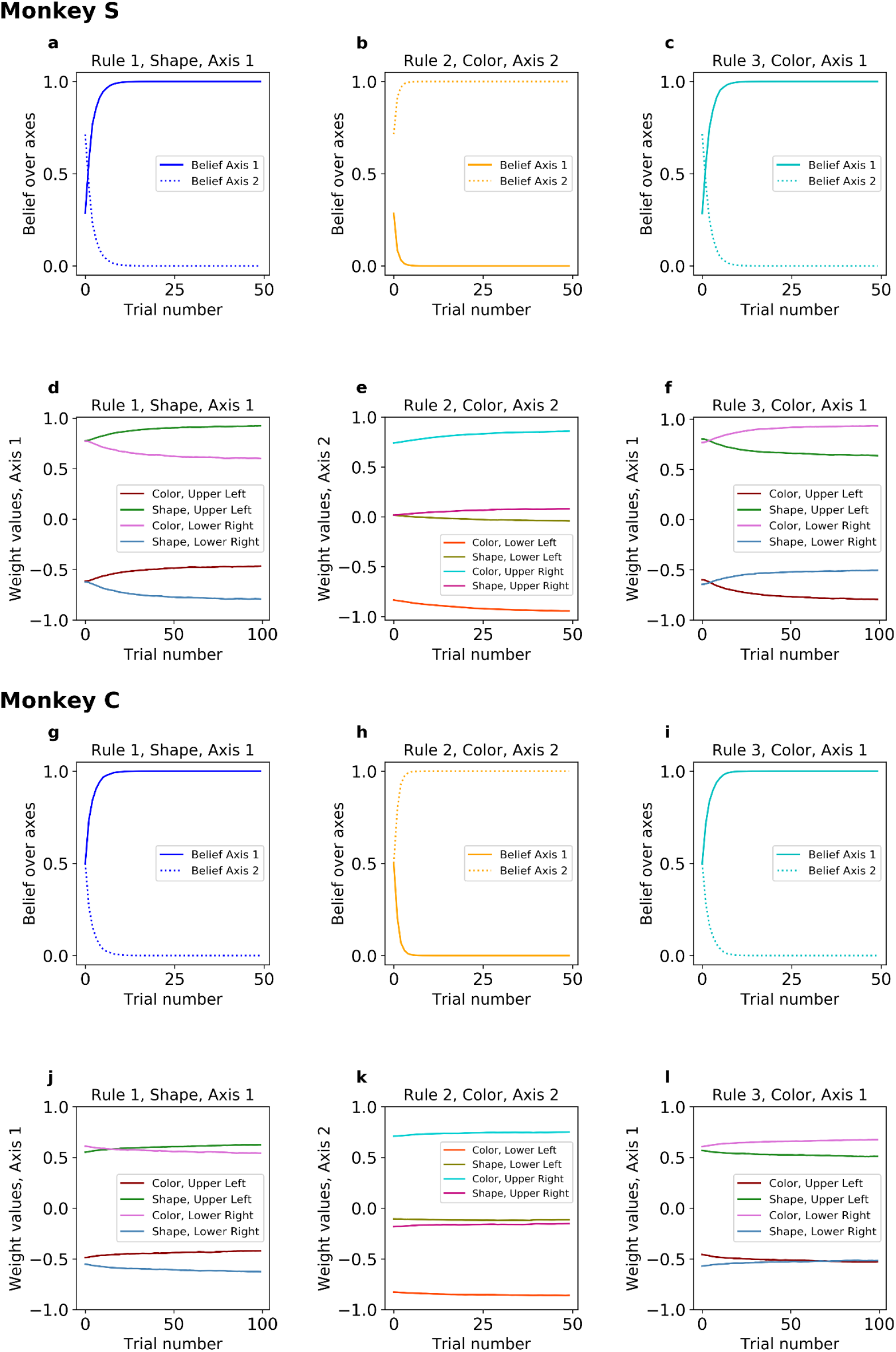
The hybrid learner. (a-c) Belief over axes when the model is fitted on Monkey S. (d-f) Feature weights values when the model is fitted on Monkey S. (g-i) Belief over axes when the model is fitted on Monkey C. (j-l) Feature weights values when the model is fitted on Monkey C.

**Figure S8:**
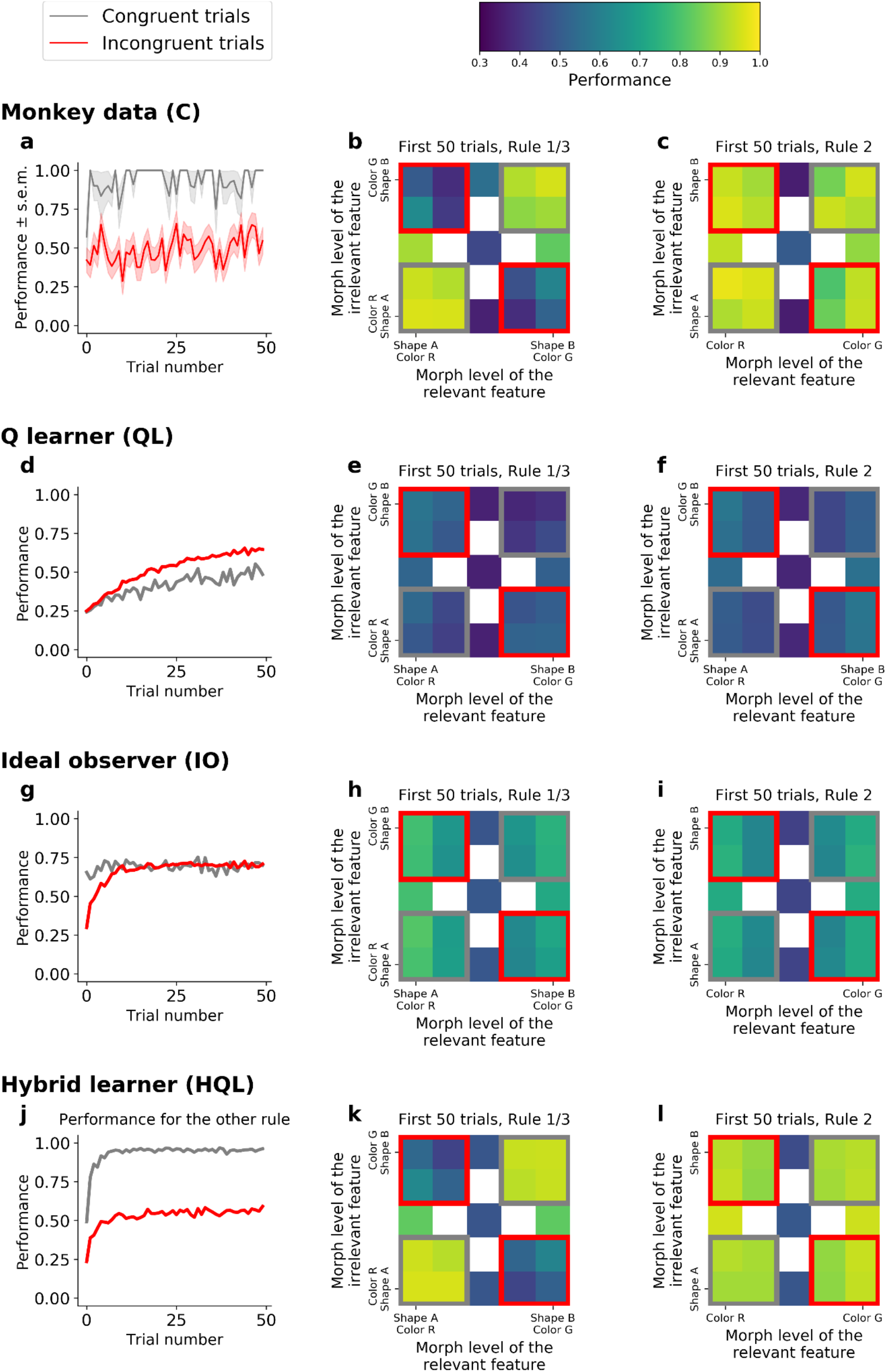
Comparison of incongruency effects in Monkey C and behavioral models (QL, IO, and HQL models). (a) Performance as a function of trial number for Rule 1 and Rule 3 (combined), for congruent and incongruent trials. (b) Performance for Rule 1 and 3 (combined, first 50 trials), as a function of the morph level for both color (relevant) and shape (irrelevant) features. Grey boxes highlight congruent stimuli, red boxes highlight incongruent stimuli. (c) Performance for Rule 2, as a function of the morph level for both color (relevant) and shape (irrelevant) features. Note the lack of an incongruency effect. (d,e,f) Same as a-c but for the QL model. (g,h,i) Same as a-c for the IO model. (j,k,l) Same as a-c but for the HQL model. <u>Statistics on Monkey C</u>: During early trials of Rules 1 and 3, the monkeys’ performance was significantly higher for congruent trials than for incongruent trials (gray vs. red squares; 93%, CI=[0.91,0.95] versus 49%, CI=[0.47,0.51] respectively; with Δ=44; Fisher test p<10^-4^). There was no difference in performance between congruent and incongruent stimuli during Rule 2 (grey vs. red squares; performance was 94%, CI=[0.91,0.95], and 91%, CI=[0.89,0.92], respectively; with Δ=2.7; Fisher test p=0.07). <u>Statistics of QL model fitted on Monkey C</u>: The model performed worse on congruent than incongruent trials in Rule 1 and Rule 3 (41% and 52%, respectively; Δ=-10; Fisher test p<10^-4^), against our behavioral observations. Furthermore, the model produced a difference in performance during Rule 2 (48% for congruent versus 54% for incongruent; Δ=-6.1; Fisher test p<10^-4^). <u>Statistics of IO model fitted on Monkey C</u>: Learning quickly reached a low asymptotic performance in Rule 1 and Rule 3, for both congruent and incongruent trials (69% and 67% respectively; Δ=2.5 only. <u>Statistics of HQL model fitted on Monkey C</u>: The model reproduced the greater performance on congruent than incongruent stimuli in Rule 1 and Rule 3 (94% and 53%, respectively; Δ=41). It also captured the absence of incongruency effect in Rule 2 (green versus red squares; 91% and 91%, respectively; Δ=0.081).

**Figure S9:**
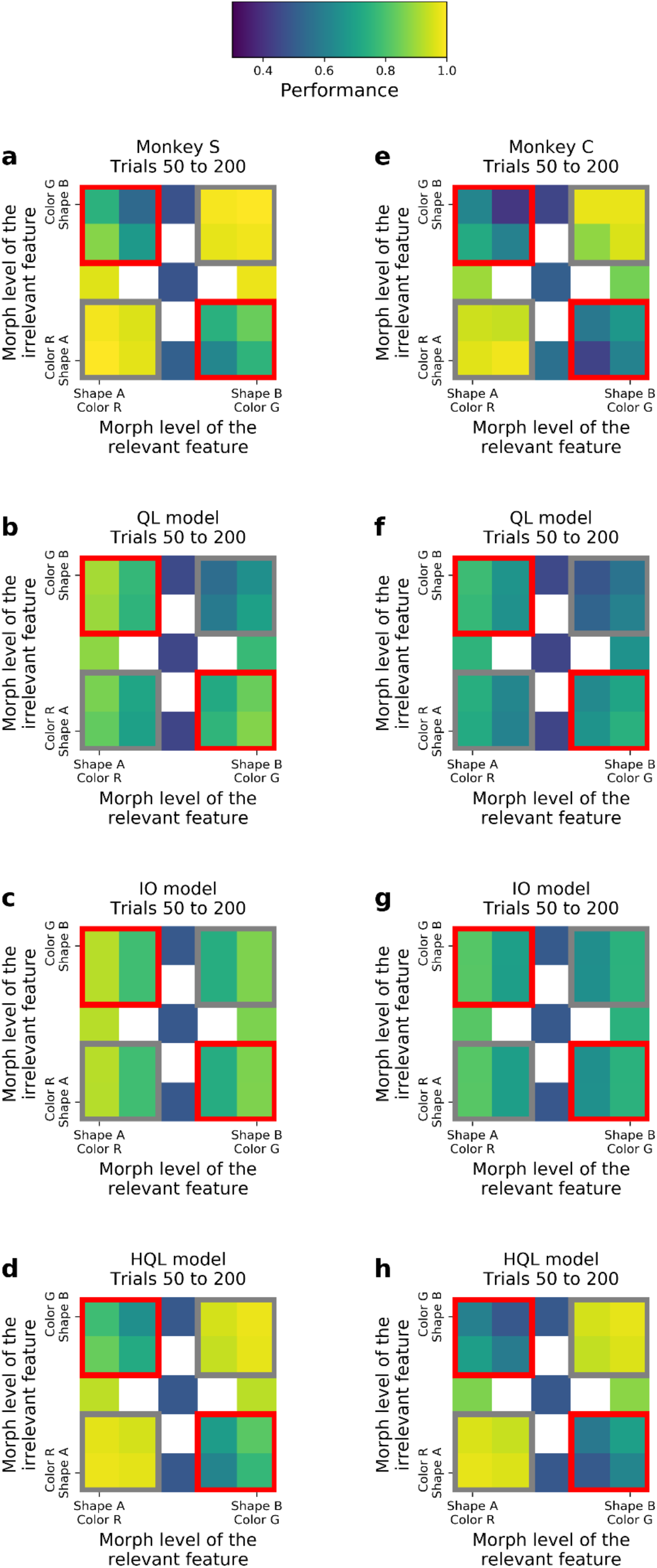
Incongruency effect for Monkey S (a), Monkey C (e), and models (respectively fitted on Monkey S: b-d; and on Monkey C: f-h), for trials 50 to 200. Each plot represents the performance for Rule 1 and Rule 3 (combined), as a function of morphs for both relevant and irrelevant features. Gray corners for congruent stimuli, red corners for incongruent stimuli.

**Figure S10:**
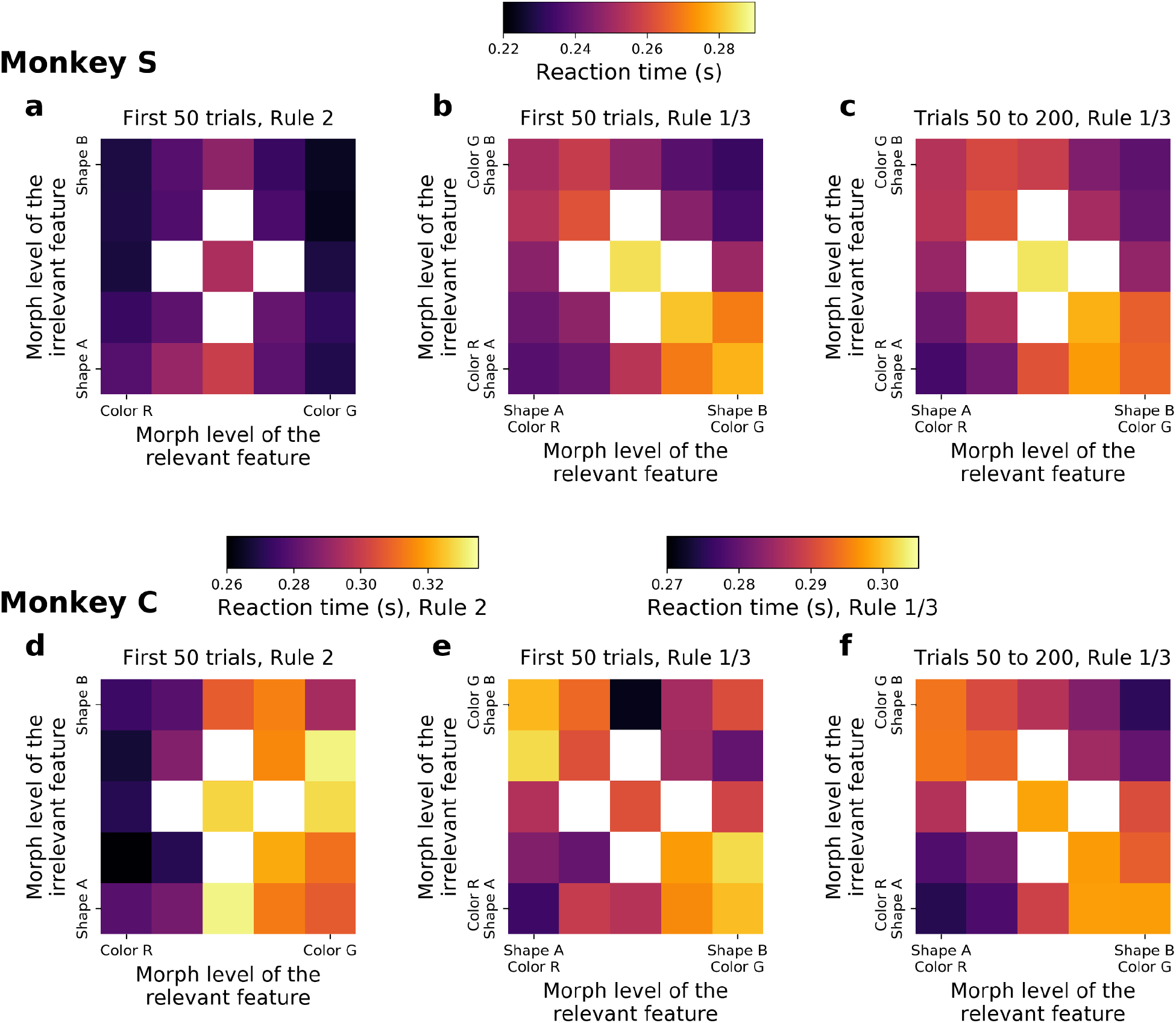
Reaction times for Rule 2 blocks, the first 50 trials of Rule 1/3 blocks, and the trials 50 to 200 of Rule 1/3 blocks as a function of the relevant and irrelevant features of the morphed stimulus presented. Top row: Monkey S, bottom row: Monkey C. <u>Statistics on Monkey C</u>: Δ(ms)=14 between incongruent and congruent, t-test p<10^-4^).

**Figure S11:**
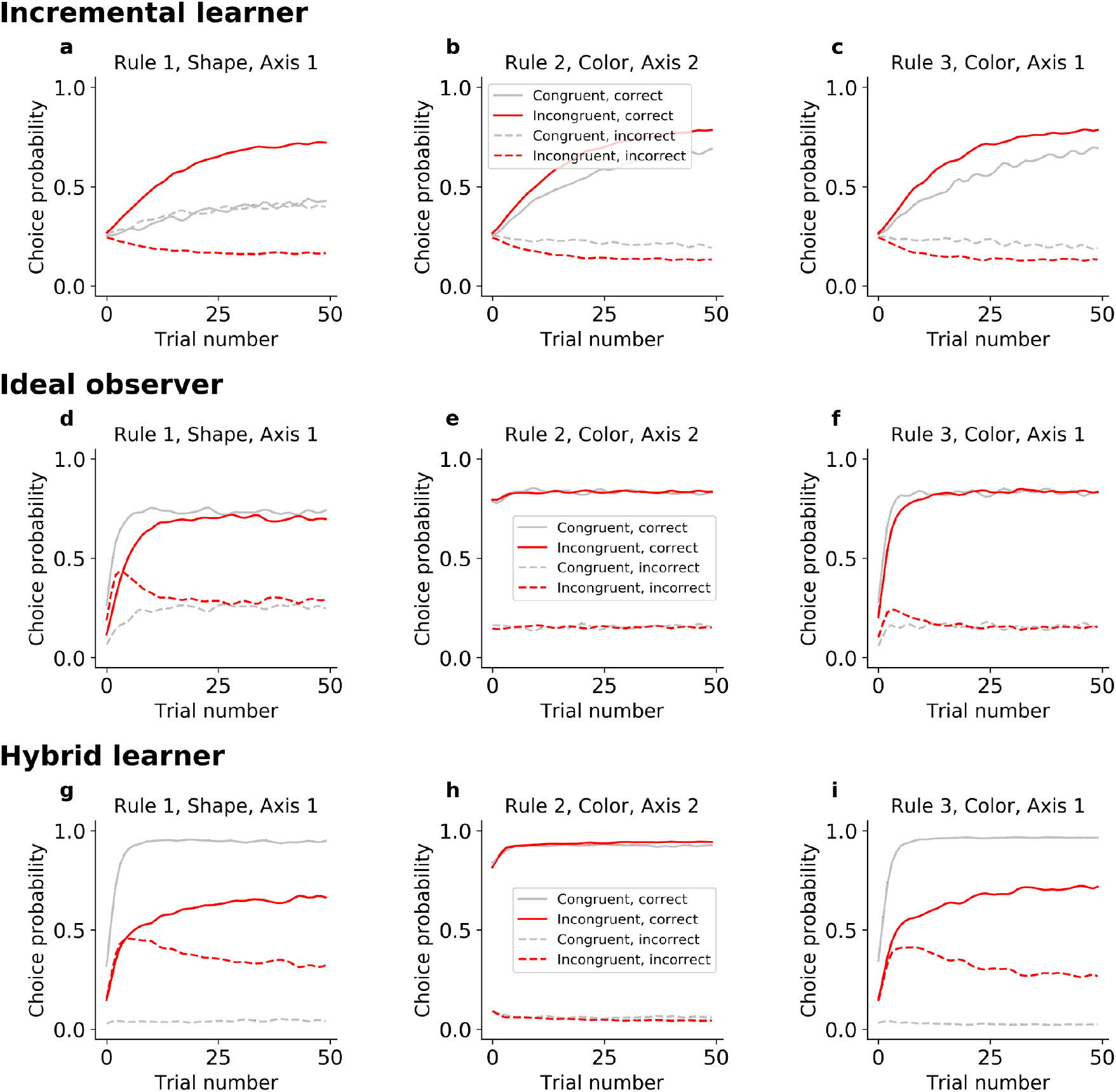
Choice probabilities for the three models fitted on Monkey S.

**Table S1:**
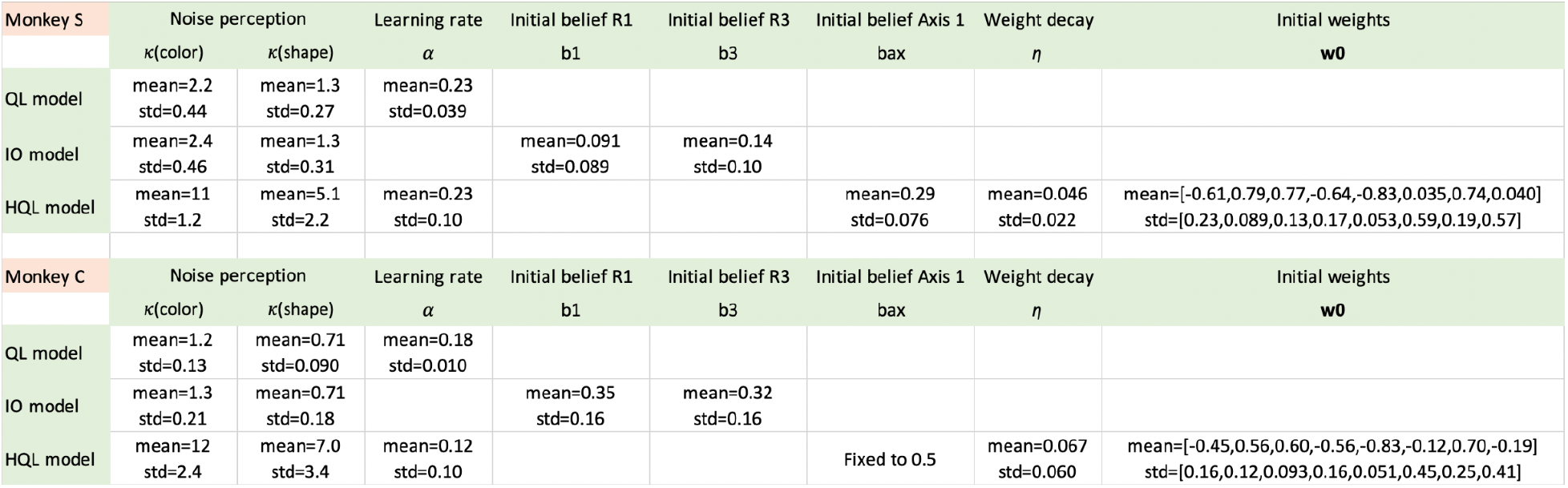
Models parameters.

| Monkey S | Noise perception |  | Learning rate | Initial belief R1 | Initial belief R3 | Initial belief Axis 1 | Weight decay | Initial weights |
| --- | --- | --- | --- | --- | --- | --- | --- | --- |
| | $\kappa(\text{color})$ | $\kappa(\text{shape})$ | $\alpha$ | b1 | b3 | bax | $\eta$ | w0 |
| QL model | mean=2.2<br>std=0.44 | mean=1.3<br>std=0.27 | mean=0.23<br>std=0.039 |  |  |  |  |  |
| IO model | mean=2.4<br>std=0.46 | mean=1.3<br>std=0.31 |  | mean=0.091<br>std=0.089 | mean=0.14<br>std=0.10 |  |  |  |
| HQL model | mean=11<br>std=1.2 | mean=5.1<br>std=2.2 | mean=0.23<br>std=0.10 |  |  | mean=0.29<br>std=0.076 | mean=0.046<br>std=0.022 | mean=[-0.61,0.79,0.77,-0.64,-0.83,0.035,0.74,0.040]<br>std=[0.23,0.089,0.13,0.17,0.053,0.59,0.19,0.57] |
| Monkey C | Noise perception |  | Learning rate | Initial belief R1 | Initial belief R3 | Initial belief Axis 1 | Weight decay | Initial weights |
| | $\kappa(\text{color})$ | $\kappa(\text{shape})$ | $\alpha$ | b1 | b3 | bax | $\eta$ | w0 |
| QL model | mean=1.2<br>std=0.13 | mean=0.71<br>std=0.090 | mean=0.18<br>std=0.010 |  |  |  |  |  |
| IO model | mean=1.3<br>std=0.21 | mean=0.71<br>std=0.18 |  | mean=0.35<br>std=0.16 | mean=0.32<br>std=0.16 |  |  |  |
| HQL model | mean=12<br>std=2.4 | mean=7.0<br>std=3.4 | mean=0.12<br>std=0.10 |  |  | Fixed to 0.5 | mean=0.067<br>std=0.060 | mean=[-0.45,0.56,0.60,-0.56,-0.83,-0.12,0.70,-0.19]<br>std=[0.16,0.12,0.093,0.16,0.051,0.45,0.25,0.41] |

## Notes

### Competing Interest Statement

The authors have declared no competing interest.

### Summary of Updates

Fig. S5 added, some explanatory text added.

